# Antigenic drift expands viral escape pathways from imprinted host humoral immunity

**DOI:** 10.1101/2024.03.20.585891

**Authors:** Daniel P. Maurer, Mya Vu, Aaron G. Schmidt

## Abstract

An initial virus exposure can imprint antibodies such that future responses to antigenically drifted strains are dependent on the identity of the imprinting strain. Subsequent exposure to antigenically distinct strains followed by affinity maturation can guide immune responses toward generation of cross-reactive antibodies. How viruses evolve in turn to escape these imprinted broad antibody responses is unclear. Here, we used clonal antibody lineages from two human donors recognizing conserved influenza virus hemagglutinin (HA) epitopes to assess viral escape potential using deep mutational scanning. We show that even though antibody affinity maturation does restrict the number of potential escape routes in the imprinting strain through repositioning the antibody variable domains, escape is still readily observed in drifted strains and attributed to epistatic networks within HA. These data explain how influenza virus continues to evolve in the human population by escaping even broad antibody responses.

## INTRODUCTION

Influenza viral escape from host humoral immunity induced by prior infection or vaccination reduces vaccine effectiveness, requiring annual updating^1^. To escape host humoral immunity, influenza acquires mutations in the surface-exposed hemagglutinin (HA) that evade antibody recognition while also maintaining functional fitness. This antigenic drift of seasonal H1N1 and H3N2 influenzas primarily occurs in the HA head domain, which engages the receptor sialic acid. Every ∼2-10 years, these mutations result in major antigenic changes in circulating influenza A viruses (IAVs) resulting in a new antigenic “cluster” that reduces the efficacy of prior immunity^2–4^. Notably, only a few amino acid positions that surround the HA receptor binding site (RBS) are largely responsible for these antigenic cluster transitions^2–6^.

Modeling IAV evolution suggests that periods of minimal antigenic change followed by antigenic cluster transitions can be explained by accumulation of “neutral” mutations that do not substantially alter antigenicity alone but influence the effects of subsequently acquired mutations^7^; mutations that precipitate antigenic cluster transitions are thus dependent on the HA identity. This “epochal” evolution model was reinforced by demonstrating that a single mutation can have no effect in one IAV strain, but after acquiring additional mutations, the same mutation results in escape from monoclonal antibodies^8^. This dependency of a mutational effect on a particular background IAV strain or the presence of other mutations (epistasis) prohibits accurate prediction of the effect of any single mutation. Previous studies have used deep mutational scanning (DMS) to understand how mutations might impact IAV escape from monoclonal antibodies, including those that target the RBS^9–12^. It was found that different IAV strains can have different barriers to escape^11^ and epistasis between mutations in the RBS can influence escape^12^. However, to examine escape mutations in the RBS, these studies used a historical IAV (WSN/1933) and primarily mouse antibodies to select for escape, which does not reflect the complex immune history that occurs in humans.

The immunological imprinting phenomenon – first termed “original antigenic sin”^13–15^ – describes that antibodies present in sera elicited by the initial IAV exposure are largely “back-boosted” and refined by subsequent exposure to drifted strains rather than generating *de novo* antibody responses. Similarly, “antigenic seniority” describes that antibody titers are higher for IAVs more antigenically similar to the imprinting strain; titers are gradually lower for IAVs that have undergone an increasing amount of antigenic drift^16^. Recent murine studies showed that the degree of back-boosting versus *de novo* responses is dependent on the antigenic distance between the HAs^17^. Larger antigenic distances between the first exposure (*i.e.,* imprinting strain) and the boosting strain (*i.e.,* drifted strain) increase the proportion of *de novo* responses and elicit serum antibodies targeting drifted epitopes due to escape from previously circulating antibodies^17,18^. In humans, IAV infection back-boosts antibodies with the greatest increase in titer towards the infecting strain, while the long-lived antibody titers remain biased towards the imprinting strain^19^. During an initial IAV infection, naive B cells engage HA and undergo somatic hypermutation and selection in germinal centers (GCs) to generate a pool of memory B cells encoding antibodies with high affinity and specificity for an epitope, a process known as affinity maturation. Subsequent exposure to antigenically drifted IAVs primarily activates these memory B cells that either re-enter GCs for further affinity maturation to the mutated epitope or directly differentiate into antibody secreting cells (*e.g.,* plasmablasts)^20,21^. By sequencing antibody variable regions from plasmablasts isolated after influenza infection or vaccination, clonally related antibodies can be identified. Phylogenetic reconstruction the unmutated common ancestor (UCA) and intermediates in affinity maturation, inferred^22–27^. Assessing the reactivity to HA from historical IAVs can identify the likely imprinting strain. Indeed, for several RBS-directed antibody lineages^22,26,27^, the UCAs only bound HAs from the antigenic cluster that circulated around the time the donor was born, but the affinity-matured antibodies could recognize strains from multiple clusters, spanning up to ∼30 years^25^. These RBS-directed lineages along with knowledge of the likely imprinting and boosting IAV strains allows interrogation of how affinity maturation influences viral escape at a conserved site and whether the barrier to escape changes in the boosting, drifted strain due to epistatic effects.

Here, we use DMS and viral escape assays in H1N1 strains from different antigenic clusters to understand viral escape in the context of imprinted antibody lineages. We find that although affinity maturation in these lineages greatly increases the barrier to escape in the imprinting strain, antigenically drifted strains readily escape these antibodies despite similar neutralization potencies. This strain dependency is due to epistasis between mutations that individually have no impact on escape, consistent with IAV evolution models. Importantly, we find that escape selections with only two RBS antibody lineages identifies positions responsible for major antigenic cluster changes in IAVs, suggesting such RBS-directed antibodies may drive viral evolution at the population level. Surprisingly, these data suggest that recalling imprinted antibody lineages through vaccination, even those that are broadly neutralizing by targeting a conserved site, expands the number of escape routes. Thus, next-generation influenza vaccines must consider how drawing from immune memory instead of generating *de novo* responses may simultaneously provide protection and increase the number escape pathways.

## RESULTS

### Imprinted antibody lineages targeting conserved HA sites

The RBS and lateral patch are conserved epitopes on the HA head and are targets of monoclonal antibodies (mAbs) that neutralize viruses from multiple antigenic clusters (**Figure 1A**). While RBS conservation is imposed by functional constraints (*e.g.,* receptor engagement), the lateral patch has no defined function^24^. Prior work isolated mAbs from human donors post-vaccination and reconstructed lineages that target the RBS or lateral patch to determine the UCAs and identify the likely imprinting strains^22–27^ (**Figure S1A**). The RBS-directed mAbs were isolated from day 7 plasmablasts of a donor born in 1989 (“1989 donor”) after trivalent vaccination in 2008, which contained the IAV H1N1 strain A/Solomon Islands/03/2006 (SI06)^23^. The lateral patch-directed mAbs were isolated from plasmablasts of a donor born in 1975 (“1975 donor”) after monovalent vaccination with a 2009 pandemic strain^24^. A common feature of these lineages is that the UCAs only recognize strains from the antigenic cluster prevalent at the time of the donor’s birth (or shortly thereafter)^24,25^. Hence, the strain bound by the UCA corresponds both to the likely first influenza exposure for that donor and to the strain that elicited the lineage. Affinity maturation of these lineages enabled the mAbs to bind later antigenic clusters, consistent with initially established and subsequently recalled imprinted antibody responses (**Figures 1B and S1A**).

**Figure 1.**
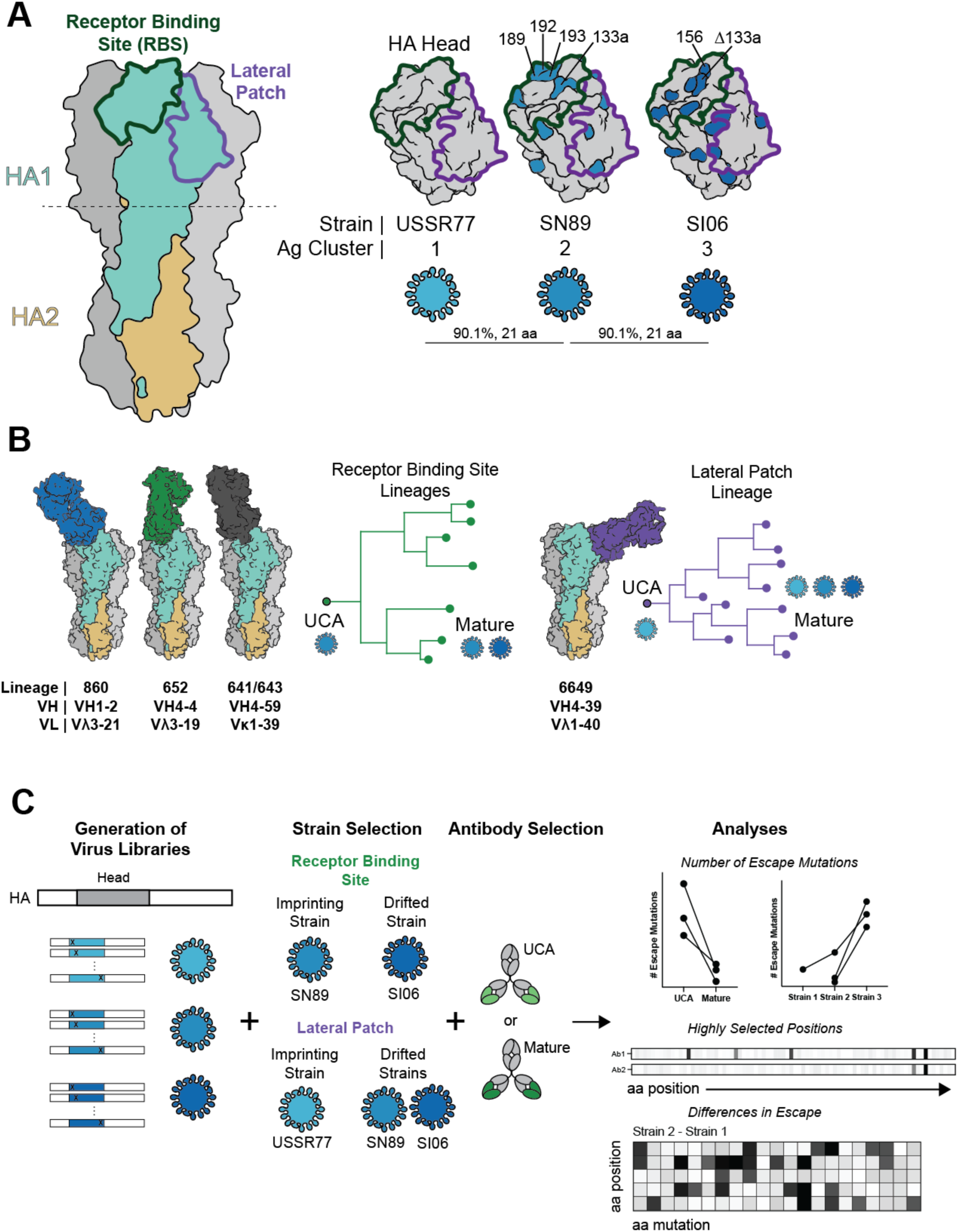
Deep viral escape from imprinted antibody lineages. **(A)** HA trimer (left) highlighting the RBS and lateral patch epitopes. Dotted line depicts head domain cutoff. HA trimer is modeled from PDB 5UGY. Mutations accrued during evolution between strains are mapped onto the head domain (right) with the associated strain and antigenic cluster. HA head is modeled from A/Massachusetts/1/1990, PDB 9AZR. Clusters are based on Bedford et al.^2^ Percent identity and number of amino acid changes in the head domain between strains are shown. **(B)** Structures from RBS or lateral patch lineages (PDB: 5UGY, 6Q0O, 4YK4, 5W6G) along with a schematic of phylogenetic reconstructions and binding to viruses colored as in A. **(C)** Schematic of approach: (1) deep mutational scanning virus libraries are generated, (2) imprinting and drifted strains are selected based on reactivity profiles of the antibody lineages, (3) the UCAs and/or mature antibodies are used to pressure for escape, (4) number of escape mutations, highly selected positions, and differences in escape are analyzed from deep sequencing data.

We chose the following lineages and IAVs for experiments to compare escape from the inferred UCAs with escape from the affinity-matured mAbs, both by the likely imprinting IAVs and by antigenically drifted strains. RBS-directed lineages 641, 643, 652, and 860, from the 1989 donor^23,25^, were presumably elicited by infection with an IAV similar to H1N1 A/Siena/10/1989 (SN89), as the UCA bound HA from this strain, but not HAs from later antigenic clusters. The affinity-matured mAbs from these lineages also bound SI06. The lateral-patch directed lineage, 6649, was probably elicited by an IAV similar to H1N1 A/USSR/90/1977 (USSR77), as the UCA bound its HA but not HAs of strains from later antigenic clusters such as SN89 or SI06. For test IAVs we chose USSR77 (imprinting strain for the 1975 donor), SN89 (imprinting strain for the 1989 donor but drifted for the 1975 donor), and SI06 (drifted for both donors) (**Figures 1B and S1A**).

### Viral library generation and experimental design

For the three IAV strains, we generated viral libraries containing all single amino acid mutations and deletions within the HA head (positions 52-263, H3 numbering) (**Figures 1C and S1B-C**). To avoid potential RBS adaptation mutations to cell lines expressing avian-type receptors, we used humanized MDCK cells, which have only α-2,6 linked sialic acid and mimic the human upper respiratory tract^28^. For each IAV strain, two independent libraries were generated and selected separately (**Figure S1D**). To promote incorporation of a single HA variant per virion, we passaged the viral libraries at low MOI (0.01). Generation and propagation of these viral libraries resulted in viral populations that depleted variants containing stop codons and deletions (**Figures S1D and S1E**). There was correlation (r ∼ 0.4-0.8) of functional scores^29^, which reflect the change in frequency of individual amino acid variants after viral passaging, between two completely independent biological replicates (**Figures S1D-S1F**).

To assess viral escape from the antibody lineages, we performed 18 deep mutational escape selections, each in biological replicate, using the UCA or mature mAbs from each lineage against the imprinting strain and against one or both drifted strains (**Figure 1C**). We analyzed the data as follows: (1) the differences between the number and identify of escape mutations in selections for both the imprinting and drifted strains, with a UCA and its affinity-matured counterpart(s); (2) the apparent barrier to escape at each amino acid position calculated by summing the individual escape effects of each amino acid mutation (**Figures 1C and S1D**).

### Viral escape in the imprinting strain is restricted by affinity-matured RBS antibodies

For the imprinting IAV strains, we tested the neutralization potency of the UCA and affinity-matured mAbs from the five lineages. Each of the affinity-matured mAbs neutralized the presumptive imprinting strain more potently than did its UCA (**Figure 2A**). The UCAs of RBS-directed lineages 860, 652, 641 neutralized SN89 virus potently enough that we could compare the identity and number of mutations allowing escape from neutralization by the UCA with the corresponding data for escape from affinity-matured mAb(s) in the same lineage (**Figures S2A, S2F, and S2K**).

**Figure 2.**
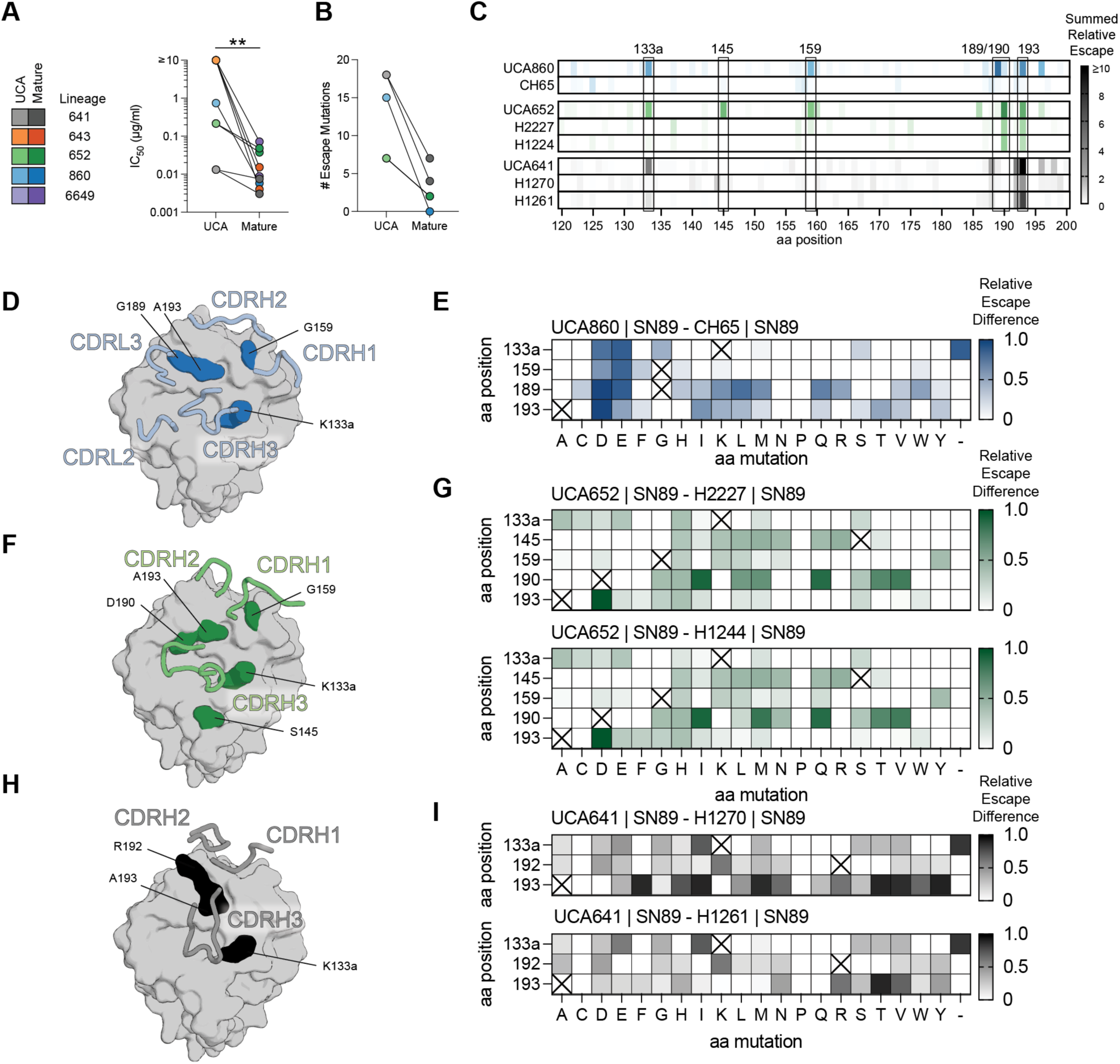
Affinity-matured RBS antibodies restrict viral escape. **(A)** Neutralization potencies between UCAs and affinity matured antibodies from each lineage. ** P < 0.005 (Wilcoxon test). **(B)** Number of escape mutations for the UCAs and mature antibodies from lineages 860, 652, and 641. Data points are colored as in A. **(C)** Relative escape values summed by amino acid position (H3 numbering). **(D,F,H)** Positions that when mutated escape the UCA more readily than the affinity-matured antibodies. Structures of antibodies bound to SI06 (PDBs 5UGY **(D)**, 6Q0O **(F)**, and 4YK4 **(H)**) are superimposed on a MA90 HA head (a strain similar to SN89). (**E,G,I**) Differences in escape between the UCA and mature antibodies from each lineage. Values are the relative escape of the UCA subtracted by the relative escape of the mature antibody. Saturated colors represent mutations that escape the UCA more readily than the mature antibodies. ‘X’ denotes the wild-type residue at each position.

We used 10x the IC99 mAb concentration for selections with the mAb present only during initial infection to select large-effect escape mutations to RBS-directed mAbs. We used differential selection^9^, which calculates the change in frequency of HA variants relative to wild-type after passaging in the presence of antibody relative to passaging without antibody, to quantify the escape effect of individual variants. We normalized each differential selection value from each replicate to the maximally selected value for comparison between selections with the expectation that maximally selected variants completely escape mAb neutralization. We defined an escape mutation as one with a normalized score of >0.5 in two biological replicates – in other words, as a mutation reproducibly positively selected with at least half the differential selection value of the most highly selected mutation (**Figures S1D, S2B, S2G, S2L and S2Q**).

We found 7-18 escape mutations from the UCAs of the RBS-directed lineages, but only 2-7 escape mutations from the affinity-matured RBS-directed mAbs. Affinity maturation thus reduced the number of escape pathways by 56-100% and eliminated escape at certain positions (**Figures 2B-2C**). For example, a total of 15 mutations at positions 133a, 159, 189, and 193 including acidic residues Glu and Asp as well as a deletion at 133a, allowed SN89 to escape neutralization by UCA860 (**Figures 2B-2E**), while affinity maturation of UCA860 to CH65 eliminated escape by viruses bearing any of these mutations, and we were unable to select for escape mutations from CH65 at any other positions (**Figures 2C and S2A-S2E**). Similarly, mutations at positions 133a, 145, 159, 190, and 193, allowed SN89 to escape neutralization by UCA652, the mutations at the first three of these positions failed to allow escape from the two affinity-matured mAbs, and there were fewer escape mutations at the remaining two positions (**Figures 2C, 2F, 2G, and S2F-S2J**). Results for lineage 641 were similar (**Figures 2C, 2H, 2I, and S2K-S2O**).

We expressed selected SN89 variants as soluble, recombinant HA trimers to determine avidities for the UCAs and affinity-matured mAbs (**Figure S2U**); We found a rank correlation (ρ = 0.78) of avidities with relative escape values (**Figures S2U and S2V**). Affinity-matured mAbs maintained wild-type avidities against mutations that otherwise reduced binding of the UCAs to undetectable levels (**Figure S2U**), and nearly all the variants we define as “escape” variants had undetectable levels of binding (**Figure S2W**). Thus, affinity maturation of RBS-directed mAbs raises the barrier to escape in the imprinting strain, by restricting the number of viral escape pathways.

### Viral escape mutations from imprinted antibody lineages are abundant in drifted strains

We compared escape of imprinting and drifted strains from neutralization by affinity-matured mAbs for RBS-directed lineages 860 and 652 as well as the lateral patch-directed lineage 6649 (**Figure 3**). In general, the mAbs neutralized the drifted strains with lower potency compared to the imprinting strain, except for the affinity-matured mAbs from lineages 860 and 652 that maintained similar neutralization potencies (<5-fold) (**Figure 3A**). Despite this modest change, we found in SI06 substantially (34-43) more mutations that escaped neutralization by the affinity-matured mAbs in lineages 860 and 652 than we found in SN89 (**Figure 3B**). In lineage 860, for which no escape mutations from CH65 in SN89 were selected, we found 34 in SI06, despite slightly less potent neutralization (4.5-fold) (**Figures 3B, 3C, and 3E**). The positions of mutations in SI06 that escaped CH65 are those of mutations in SN89 that escaped UCA860 with more possible substitutions (**Figures 3C, 2C, and S3A-S3F**). Similarly in lineage 652, affinity-matured mAbs H2227 and H1244 neutralized SN89 and SI06 equivalently, but 35-45 mutations allowed escape in SI06 compared with only 2 in SN89 (**Figures 3B, 3C, and 3F**). As we observed for lineage 860, positions in SI06 conferring escape from H2227 and H1244 in SI06 are the same as those in SN89 conferring escape from UCA652 (**Figures 3C, 3C, and S3G-S3L**). In the lateral patch-directed lineage, for which we had two drifted strains, we observed a progressively lower barrier to escape in IAVs as the antigenic distance from the imprinting strain increased (**Figures 3B, 3D, 3G, and 3H**). Not only did more escape mutations appear, but more mutations at additional positions (*e.g.,* positions 128, 166, and 171) escaped mAb neutralization (**Figure 3D, 3G, 3H, and S3M-S3R**).

**Figure 3.**
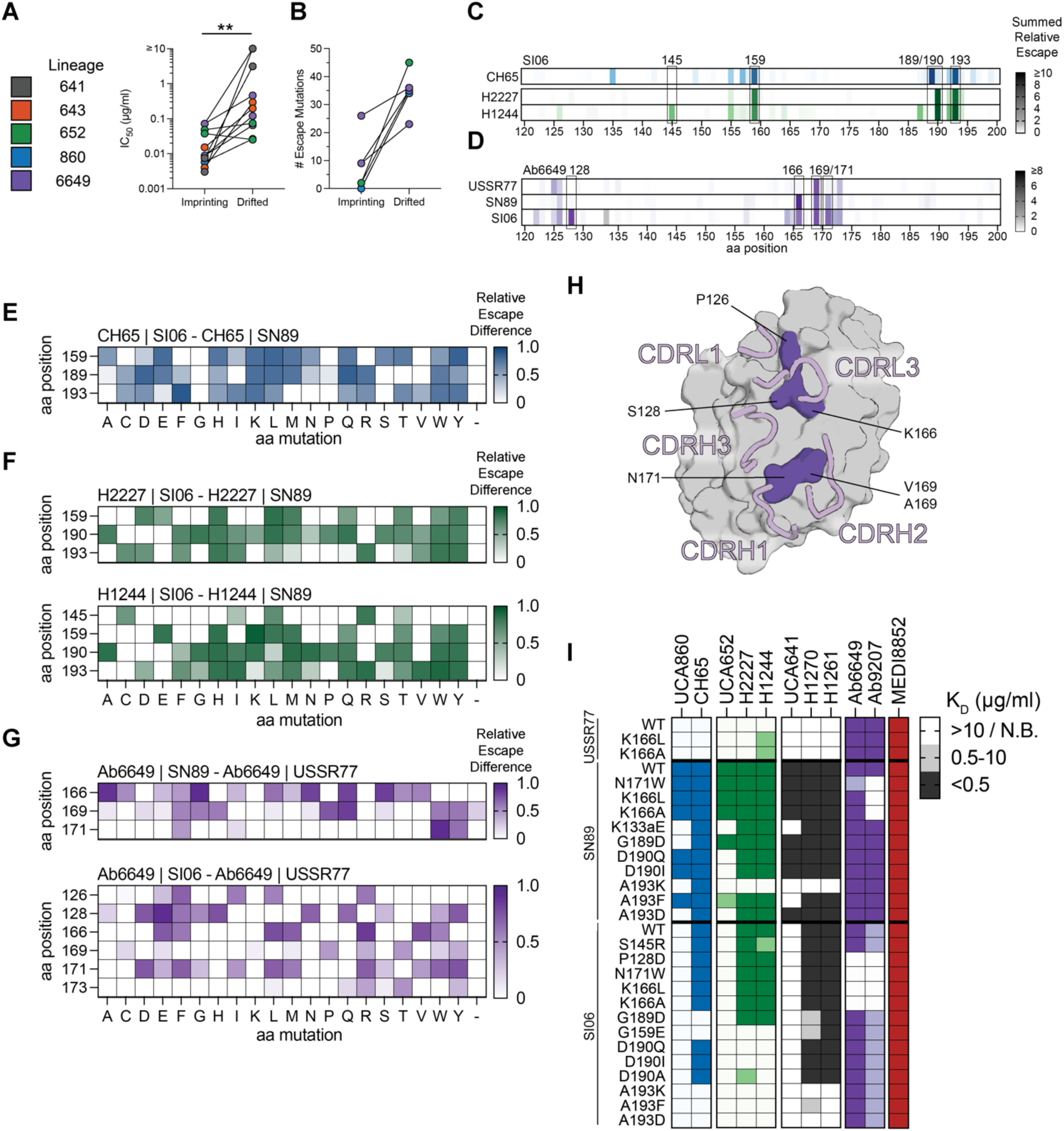
Antigenic drift expands viral escape pathways from imprinted antibody lineages. **(A)** Neutralization potencies between the imprinting strain and the drifted strain. ** P < 0.005 (Wilcoxon test). **(B)** Number of escape mutations between the imprinting and drifted strains. Data points are colored as in A. For RBS lineages, imprinting strain is SN89, drifted strain is SI06. For the lateral patch lineage (purple), imprinting strain is USSR77, drifted strains are SN89 and SI06. **(C)** Relative escape values in SI06 summed by aa position (H3 numbering) against RBS antibodies. **(D)** Relative escape values summed by aa position for Ab6649 against USSR77, SN89, and SI06. **(E-G)** Differences in escape between the imprinting and drifted strains. Values are the relative escape of the drifted strain subtracted by the relative escape of the imprinting strain. Saturated colors represent mutations that escape the antibody in the drifted strain more readily than in the imprinting strain. **(H)** Positions that when mutated escape the antibody in the drifted strain more readily than in the imprinting strain. Structure is from PDB 5W6G. **(I)** Avidities determined by ELISA of variants in each background with four lineages. MEDI8852 targets the stem region and is used as a positive control.

We next assessed avidities for a panel of 28 recombinant HAs with different observed escape mutations in different IAV backgrounds (**Figure 3I, S3S, and S3T**). Mutations G189D and A193F/K/D had no effect on CH65 affinity for the imprinting strain (SN89), but reduced binding to undetectable levels in the drifted strain (SI06) (**Figure 3I**). Similarly, mutations D190Q/I and A193F/D eliminated detectable binding of H2227 and H1244 only in the context of the drifted strain (SI06) (**Figure 3I**). Although we were unable to select for SI06 escape variants in this assay for lineage 641 due to low neutralization potency, avidity of some variants selected by other RBS antibodies (*e.g.,* A193F/D) had no effect in SN89 on inhibition of lineage 641 affinity-matured mAbs, but a very strong effect in SI06 (**Figure 3I**).

The lateral patch-directed lineage had distinct differences between clonally related members Ab6649 and Ab9207 (**Figures 3I and S3M-S3R**). Both mAbs bound HAs bearing mutations K166L/A in the imprinting strain (USSR77), but these mutations substantially reduced Ab9207 avidity for SN89 with minimal effect on Ab6649 binding. In the further drifted strain, SI06, the same mutations now eliminate detectable Ab6649 avidity. Mutation Thr173 escaped neutralization by both Ab6649 and Ab9207 (**Figure S3Q**); it introduces a predicted N-linked glycosylation site (at Asn171); a previously identified natural escape mutation in this epitope introduces a glycosylation site at position 165^24^.

Members of lineages 652 and 641 also selected for escape mutations that were not shared between affinity-matured mAbs of the same lineage. In lineage 652, the mutation S145R in SI06 escaped H1244 and substantially reduced avidity, but this mutation had no impact on the H2227 avidity (**Figure 3I and S3J-K**). In lineage 641, mutations G189D and G159E eliminated detectable binding avidity for H1270, but not H1261 (**Figure 3I**). These data show that there are more escape pathways possible from antibody clonal lineages in drifted IAV strains than in the eliciting IAV strain; the result is a lower barrier to escape from an oligoclonal antibody response for drifted strains when selected with affinity-matured members of antibody lineages originally elicited by an earlier strain (**Figure S4**).

### Antibody tilting eliminates escape potential

To understand how an affinity-matured mAb restricts viral escape, we determined a crystal structure of UCA860 bound to MA90 (an SN89-like imprinting strain) and two crystal structures of CH67 bound to MA90-G189E, which escapes UCA860 but not affinity-matured CH65 or CH67 (**Figure 4A and Table S1**). Superimposing UCA860:MA90 and CH67:G189E complexes shows that the escape mutation G189E directly clashes with the CDRL3 of the UCA860 (**Figures 4B, C**). In the CH67:G189E complexes, the variable domains “tilt” towards the 130- and 150-loops of the HA by ∼10°, allowing the CDRL3 to shift away alleviating the steric clash (**Figures 4B, C**). Prior work identified that somatic mutations N52H and G31D, which occurred early during affinity maturation^25,27^, compensate for the escape mutation G189E^30^. In the CH67:G189E structure, these residues engage the 150-loop of HA, providing new stabilizing contacts to tilt away from the distant clash with G189E (**Figure 4C**). Despite this tilting, the CDRH3 with the critical dipeptide that mimics sialic acid is similarly positioned within the RBS, providing an anchor point. Notably, the two CH67:G189E structures appear to engage the G189E escape variant through different conformations (**Figure S5**). We observed that the CDRL2, for which there is density present in only one of the structures, forms a salt bridge with the CDRL1 through the somatic mutation S29R, altering its configuration (**Figure S5**). This different CDRL1 configuration allows Y108 to adopt an alternate rotamer that subsequently allows F110 in the CDRH3 and W90 in the CDRL3 to also undergo ring “flips”. Ultimately, the flip of W90 provides space for the CDRL3 to adopt an altered conformation that avoids clashing with the G189E mutation. These data show that somatic mutations can enable new stabilizing contacts distant from the escape mutation that prevent escape by allowing the antibody to tilt away from the site.

**Figure 4.**
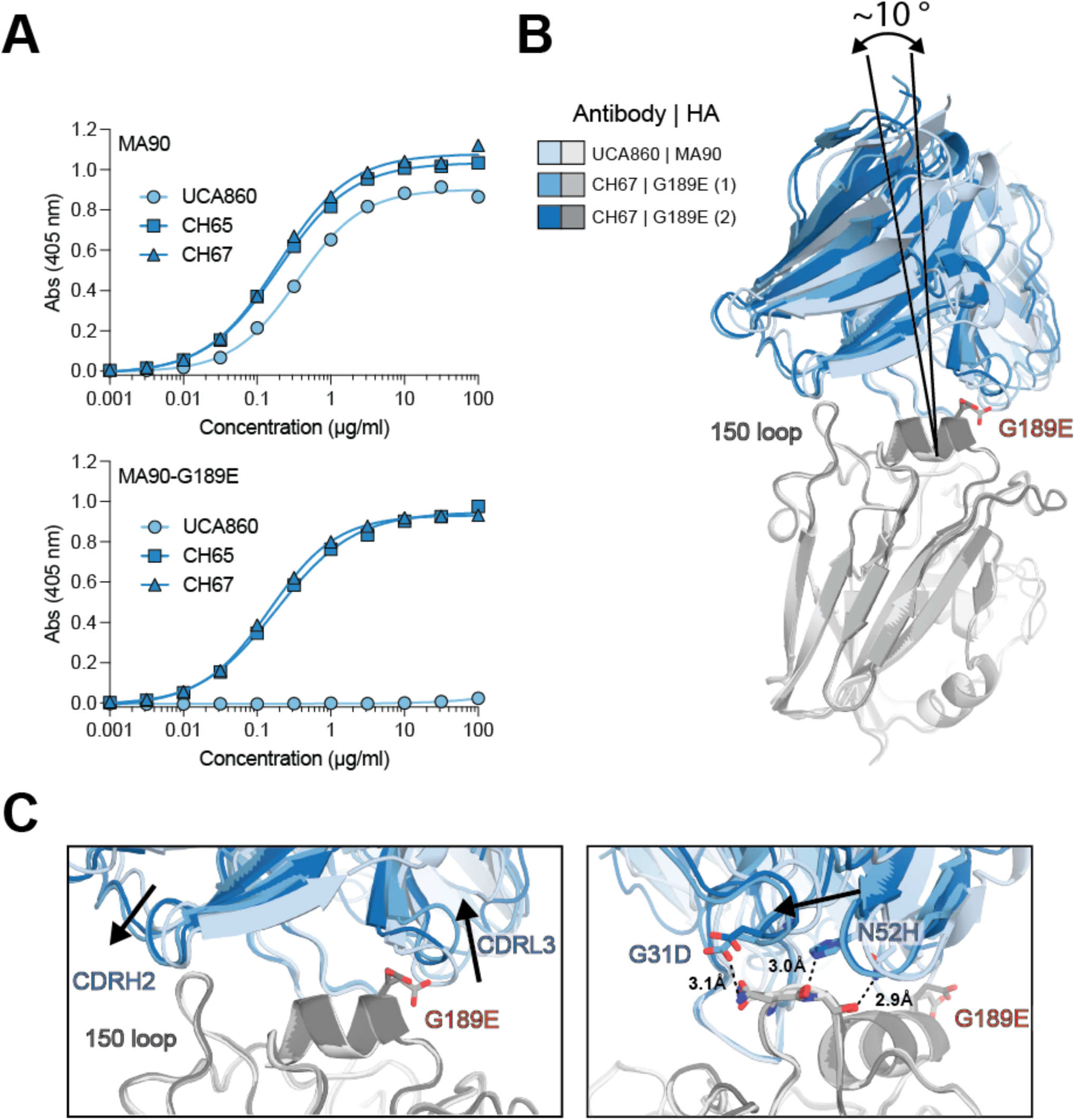
Affinity maturation alleviates clashing with an escape mutation by antibody tilting. **(A)** Reactivity of lineage 860 members to MA90 (top) and MA90-G189E (bottom). **(B)** Superimposition of UCA860:MA90 and CH67:G189E structures. Tilting angle is calculated from the tip of the CDRH3. **(C)** Zoom in of tilting motion and details of somatic mutations G31D and N52H alleviating the clash. Arrows indicate positional differences between the CDR loops of the UCA and CH67.

### Epistasis increases the number of escape pathways from an RBS-directed antibody in the drifted strain

Epistasis can account for the changes in escape and avidity by a single amino acid mutation in different background strains. To understand how the observed non-additive effects of multiple mutations might explain these changes, we compared the sequences and structures of the imprinting strain (SN89 or MA90) with that of the drifted strain (SI06). Two mutations in the RBS occurred during natural evolution from SN89/MA90 to SI06: a deletion of Lys at position 133a (ΔK133a) and E156G. In the UCA860:MA90 structure, the CDRH3 residues E102 and P103 interact with K133a and residues Y109 and Y111 interact with E156 (**Figure 5A**); thus, evolution of SN89/MA90 to SI06 with mutations ΔK133a and E156G removes these critical contacts within the antigen combining site (**Figure 5B**). These interactions suggest that binding of CH65 and CH67 to HA bearing G189D (which eliminates detectable avidity only in SI06) is contingent on the contacts with either K133a and/or E156. We generated SI06 containing the escape mutation G189D with or without reverted mutations +K133a and/or G156E, and measured avidities by ELISA (**Figure 5C**). Only when both residues were present did lineage 860 members bind this variant with wild-type avidity (**Figure 5C**). Thus, epistasis (non-additive effects of G189D combined with mutations at 133a or 156) enable escape when no single amino acid in the imprinting strain escapes CH65.

**Figure 5.**
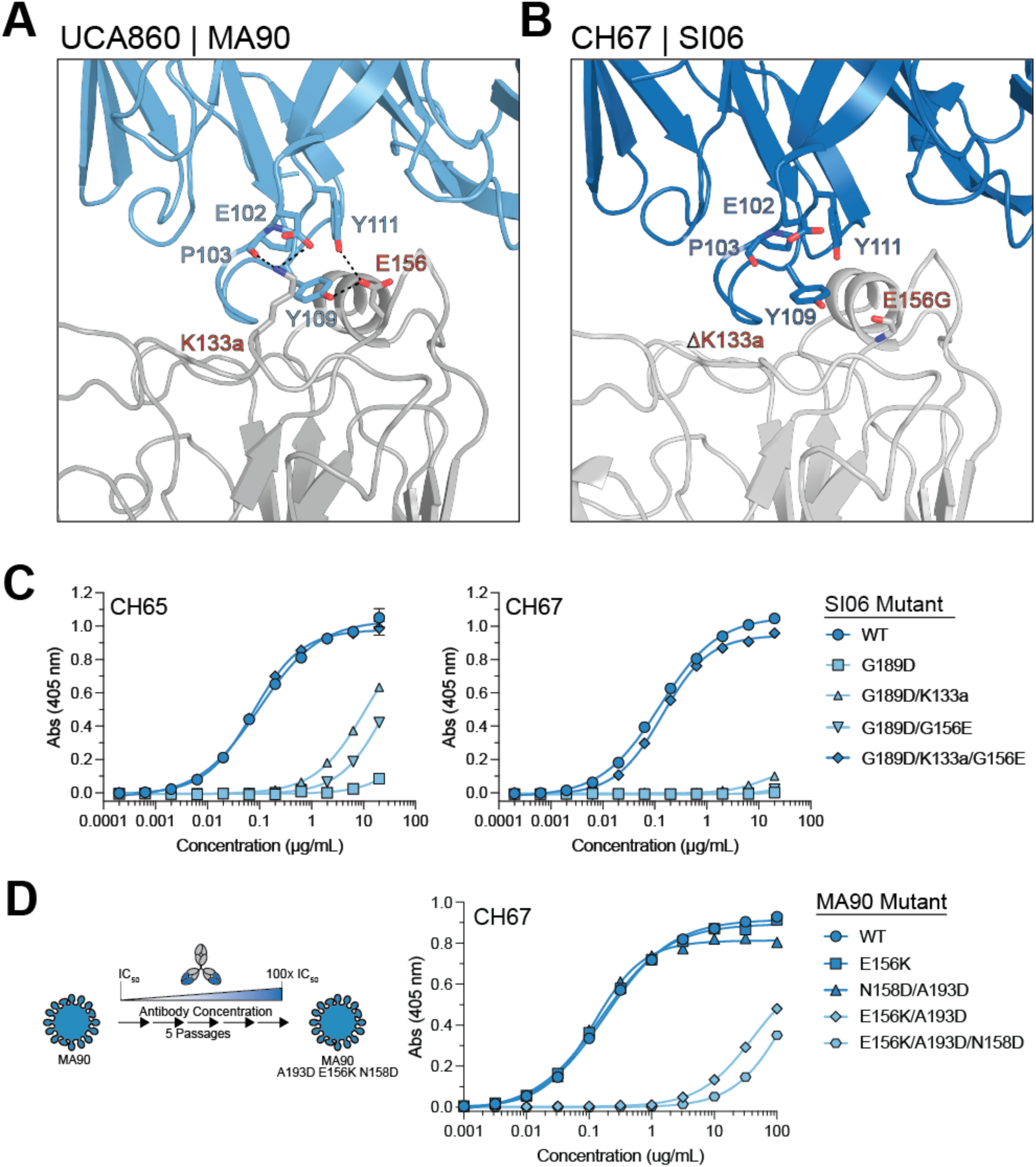
Antigenic drift enables viral escape through epistasis. **(A)** Structure of UCA860 bound to MA90, highlighting contacts between the antibody and K133a/E156 in HA. **(B)** Structure of CH67 bound to SI06 with contacts shown in A now absent. **(C)** ELISA data of CH65 and CH67 binding to SI06 variants. **(D)** MA90 virus was passaged in the presence of CH67 five times up 100x the IC50 value (left). ELISA binding data of CH67 binding to MA90 variants (right).

We passaged MA90 virus with increasing concentrations of CH67 to allow for multiple mutations to accrue. The resulting virus had a mixed population of E156K, A193D, and N158D. We assayed avidity to recombinant HA variants alone or in combination. Alone, the mutations did not affect binding of CH67, but when E156K and A193D were combined, we found a ∼200-fold avidity loss, with further loss by introduction of N158D (**Figure 5D**). Thus, there appears to be an epistatic effect of mutations at positions 156 and 193, which together enable viral escape from CH67 while neither does so on its own.

## DISCUSSION

In this study we find that affinity maturation reduces the possible number of escape mutations, but antigenic drift enables multiple escape pathways from imprinted antibody lineages targeting the RBS and lateral patch (**Figure 6**). The increase in escape mutations in drifted strains is not restricted to a particular lineage or epitope, as we find similar conclusions regarding escape from lineages that target distinct epitopes, use different variable germline genes, and are isolated from different donors. Moreover, we find that boosting imprinted antibody lineages can yield potently neutralizing antibodies in agreement with a study demonstrating antibodies with original antigenic sin phenotypes can contribute to protective responses^31^. Our data, however, suggests that boosting imprinted antibodies increases the number of escape routes in a hierarchical manner, consistent with antigenic seniority.

**Figure 6.**
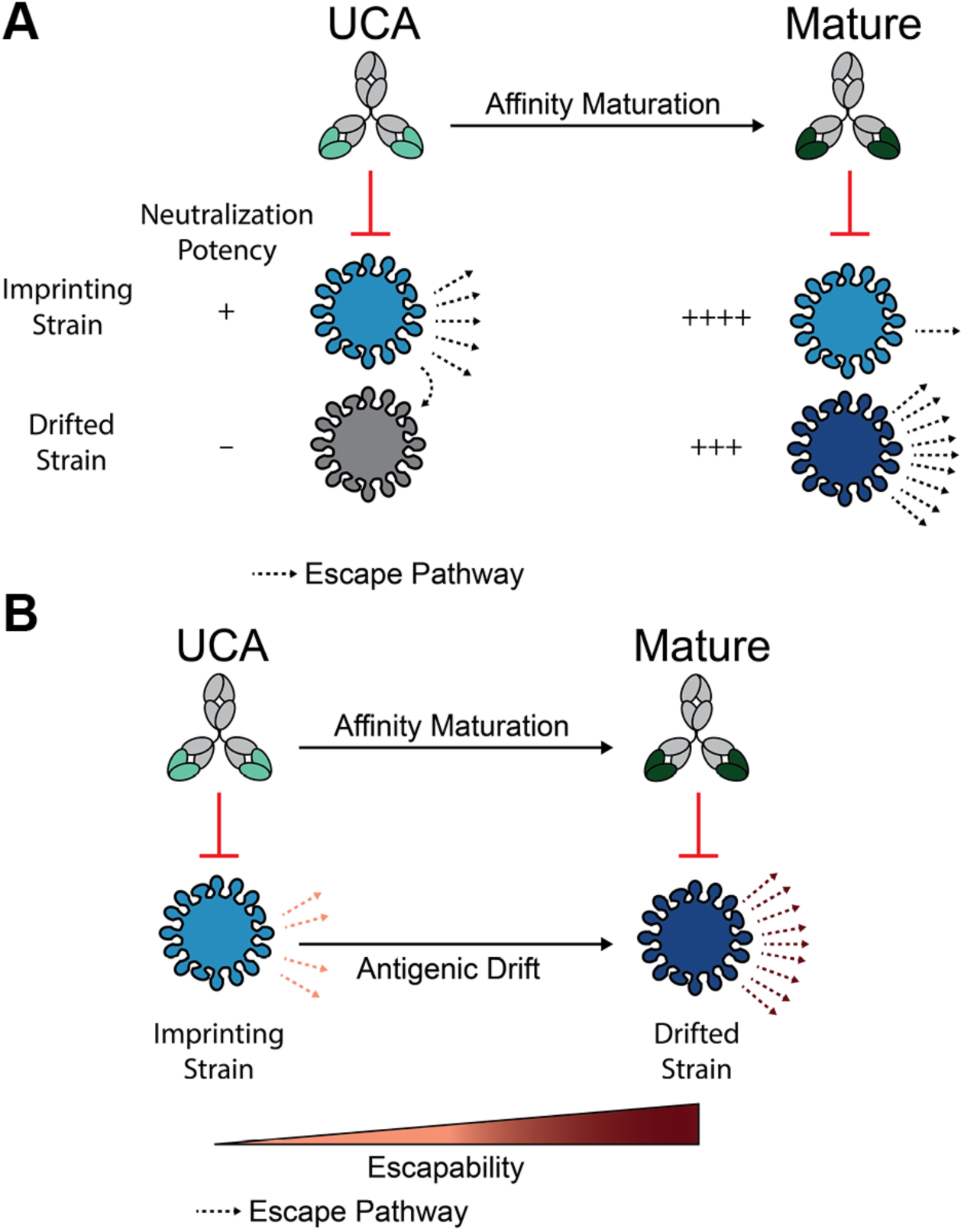
Viral escape from imprinted antibody lineages. **(A)** Affinity maturation restricts the number of viral escape pathways in the imprinting strain while also gaining neutralization potency to a drifted strain. The drifted strain has more escape pathways than the imprinting strain. **(B)** Antigenic drift affords the virus additional escape pathways from an imprinted antibody lineage.

The influenza-specific serum repertoire after vaccination is oligoclonal with only ∼1-10 clonotypes representing most of the serum^20,32^. We tested antibodies from four RBS lineages isolated from a single individual^26^, suggesting escape from an oligoclonal RBS antibody response may look similar to what we report here. The positions selected from only two RBS-directed lineages overlap with the residues known to drive antigenic change in IAVs^3,5,6^ (133a, 145, 156, 159, 189, 190, 193), and mutations at these positions almost exclusively account for escape from these RBS-directed lineages. These data suggest that recalling imprinted antibody lineages targeting the RBS may drive natural evolution more than previously thought and escape from such antibodies may be able to predict antigenic clusters that are yet to emerge.

The HA stem is currently a major target for so-called “universal” vaccine candidates, despite the clinical failure of prophylactic administration of the most broadly reactive and further engineered stem-targeting antibody (NCT05567783). The UCAs of two broadly reactive stem-directed antibodies, CR9114 and FI6, only bound group 1 HAs and acquired group 2 HA reactivity through affinity maturation^33,34^, similar to our observations with the lineages described here. When assaying for escape using deep mutational scanning in H1N1 or H3N2 viruses, escape was difficult for H1N1 (group 1), but readily occurred for H3N2 (group 2)^11^. Thus, even for anti-stem antibodies, the escape barrier appears to be high in the imprinting strain, but not the strain the lineage gains reactivity to through affinity maturation. This is not unique to IAV as epistatic effects in the SARS-CoV-2 receptor binding domain also modulate whether amino acid mutations escape neutralization^35^. These observations suggest that recalling imprinting lineages with subsequent affinity maturation may universally increase the number of mutations accessible for viral escape. The changing effect of single amino acid mutations in one IAV background compared to another highlights how significantly the background context influences the barrier to escape. For example, CH65 has no single amino acid escape mutations in SN89 but acquiring only one or two mutation(s) (ΔK133a and/or E156G) that by themselves has no effect enables a change from no escape mutations to a possible 35 mutations that can escape recognition. These data are consistent with the “neutral network” model of influenza epochal evolution^7^ that suggests acquiring mutations that themselves do not result in antigenic change can facilitate large antigenic change upon acquiring a second mutation.

This “switch-like” regulation from few to many escape pathways by minimal antigenic drift may inhibit the ability to predict how a virus may evolve in response to novel immune pressures elicited by next-generation vaccines or monoclonal antibody therapies. Computational methods aimed at predicting viral evolution require knowledge of existing evolution (*i.e.,* naturally occurring sequences)^36,37^, but the goal of many vaccination approaches is to change the dominant immune pressure by focusing antibody responses to conserved epitopes. Here, however, we show that escape from RBS antibodies select similar positions as those currently acquired on a global level, suggesting existing sequence information may provide valuable insight into escape from next-generation vaccines focusing to the RBS.

Monoclonal antibodies derived from recall frequently have the highest affinity for the imprinting strain^31,38–41^, as is the case for lineage 860^30^. Consistently, despite similar neutralization potencies between SN89 and SI06, affinity-matured mAbs from imprinted lineages 860 and 652 retain the highest barrier to escape in the imprinting strain. These data suggest that affinity maturation in these lineages biases both affinity and the barrier to escape towards the imprinting strain, “committing” antibody recognition to a historical HA. However, because affinity maturation to antigenically drifted variants has the potential to increase the barrier to escape, next-generation vaccine design should focus on immunizing with *a priori* predicted or experimentally determined antigenic clusters rather than attempting to “guide” affinity maturation towards conserved epitopes^42^ through sequential vaccination using historical strains. Although the affinity-matured antibodies more potently neutralize the drifted strains than the UCAs neutralize the imprinting strain, there are fewer escape mutations in the imprinting strain that evade the UCAs compared to number of mutations in the drifted strain that escape the mature antibodies. This finding suggests an alternative strategy is to focus on generating *de novo* antibody responses through vaccination, though additional work is required to investigate the barrier to escape from *de novo* versus recall antibody responses.

### Limitations of study

Although the levels of somatic hypermutation in RBS antibodies described here suggest multiple rounds of affinity maturation^25^, it is unknown whether these antibodies indeed underwent affinity maturation towards SI06 or a SI06-like strain. Vaccination with SI06 likely recalled memory B cells directly to plasmablasts^43^. Therefore, it may be possible that further affinity maturation additionally restricts potential escape mutations in SI06. In addition to epistasis within HA, intergenic epistasis between HA and NA also influences the accessible escape mutations in HA^44,45^. For each virus library, we used the matched NA from that strain. Thus, differences in NA may also play a role in the mutations that were selected. We used mAbs isolated from day 7-8 plasmablasts of two donors. Although these mAbs contributed to the short-lived plasmablast response, it remains to be shown whether this effect is similar in the long-lived plasma cell compartment, which would additionally contain *de novo* antibody responses.

## ACKNOWLEDGMENTS

We acknowledge support from R01 AI146779 and P01 AI089618. This research has been funded in whole or part with federal funds under a contract from the National Institute of Allergy and Infectious Diseases, NIH contract 75N93019C00050 (AGS). We thank Jesse Bloom for providing the pHW-PR8 and pHW2000 plasmids used in this work. We acknowledge the Bauer Core for PacBio library preparation and sequencing in addition to Illumina sequencing. This work is based upon research conducted at the Northeastern Collaborative Access Team beamlines, which are funded by the National Institute of General Medical Sciences from the National Institutes of Health (P30 GM124165). The Eiger 16M detector on 24-ID-E is funded by a NIH-ORIP HEI grant (S10OD021527). This research used resources of the Advanced Photon Source, a U.S. Department of Energy (DOE) Office of Science User Facility operated for the DOE Office of Science by Argonne National Laboratory under Contract No. DE-AC02-06CH11357. We thank the beamline staff at NE-CAT for assistance in data collection. We thank Angela Phillips for helpful discussions on analyses and Emerson Glassey for assistance with coding. We thank Steve Harrison for feedback on the manuscript.

## AUTHOR INFORMATION

### Author Contributions

D.P.M., A.G.S. designed research; D.P.M. and M.V. performed research; D.P.M., A.G.S. analyzed data; D.P.M., A.G.S. wrote the paper. D.P.M., M.V., A.G.S. edited and commented on the paper.

## MATERIALS AND METHODS

### Cell lines and media

293-F (ThermoFisher, #R79007) and Expi293F cells (ThermoFisher, #A14527) used for recombinant protein production were maintained in FreeStyle (ThermoFisher, #12338026) or Expi293 media (ThermoFisher, #A1435101) media, respectively, according to the manufacturer’s suggestions. HEK293T (ATCC, #CRL-3216) and humanized MDCK (hCK)^28^ cells were maintained in DMEM (ThermoFisher, #11965092) supplemented with 10% heat-inactivated fetal bovine serum (Peak Serum) and 100 units of penicillin-streptomycin (ThermoFisher, #15140122). Media used during and after influenza virus infection, “flu media”, was comprised of Opti-MEM (ThermoFisher, #31985070), 0.3% bovine serum albumin (Sigma, #A9418), 100 µg/ml calcium chloride, 100 units of penicillin-streptomycin, and 0.01% heat-inactivated fetal bovine serum. Flu media used during infection and post-infection was additionally supplemented with 1 µg/ml TPCK-treated trypsin (Sigma, #T1426).

### Recombinant protein production

Recombinant antibodies and hemagglutinin were produced following similar protocols. The variable heavy and light regions of the antibodies were cloned into a pVRC8400 backbone (VRC, NIAID, Bethesda, MD) containing human IgG CH1-CH3. HA ectodomain (or head-only) sequences (see below for accession numbers) were cloned into a pVRC8400 backbone containing a C-terminal 3C cleavage site followed by a His tag and AviTag. The full-length ectodomain recombinant HA trimers included three mutations, H26W/K51I/E103I, shown to increase expression and stability of pre-fusion HA and express full-length trimers without the requirement of a trimerization tag^46^. Antibodies and HAs were produced by production in 293F cells or Expi-293F cells, respectively, using PEI (3:1 ratio) for transfection of 293F cells and ExpiFectamine (ThermoFisher, #A14525) for transfection of Expi-293F cells according to the manufacturer’s suggestion. After 5-7 days, supernatant was clarified by centrifugation. TALON metal affinity resin (Takara, #635653, 0.5-1:1 v/v ratio) was washed with PBS and added to the supernatant. The mixture was incubated at 4 °C, rolling, for 30-60 minutes before gravity filtration (Bio-Rad, #7321011). The resin was washed with ∼5-10 bed volumes of PBS before adding 20 mls of PBS containing 200 mM imidazole (pH ∼7.4; elution buffer). After 5 minutes of incubation with elution buffer, the cap was removed and the elution was collected in protein concentrators (ThermoFisher, #88531, 30K MWCO). The elution was concentrated under 0.5 ml and filtered by spin filtration (0.22 µm) before purification over an Superdex 200 Increase 10/300 GL on an AKTA Pure system (Cytiva). For x-ray crystallography, recombinant HA heads were produced in 293F cells for the UCA860:MA90 complex and in 293 GnTI knockout cells for the CH67:MA90-G189E complexes.

### Virus library design

Viral libraries were designed to cover all single amino acid mutations in the HA head domain (positions 53-263, H3 numbering) of three H1N1 virus strains: A/USSR/90/1977 (USSR77, EPI230599), A/Siena/10/1989 (SN89, EPI171157), A/Solomon Islands/03/2006 (SI06, EPI155197). A known egg adaptation mutation in SI06 (Q226R) was reverted to Q226.

Rather than sampling all codon mutations, we used the most frequently used codons from influenza A viruses from > 40,000 coding sequences in the Kazusa database^47^ (**Table S2**). This reduced nucleotide diversity of the library and ensured comparison of codon changes between strains was not due to effects caused by specific codon usage. Because cloning into the reverse genetics vector uses the BsmBI restriction enzyme (see below), in the rare cases that the selected codons introduced a new BsmBI site, we modified the surrounding nucleotides such that introduction of a codon mutation never introduced a new BsmBI site. Additionally, these codons did not introduce SalI restriction sites, which was used to linearize the cloned plasmid libraries prior to long-read sequencing (see below). In addition to non-synonymous mutations, we also aimed to introduce synonymous and stop codons at a frequency of ∼1-2% of the library to provide an approximate measure of fitness. Every fourth position was mutated to a stop codon in the head and nonsynonymous codons were added wherever wild-type did not match the selected codon usage from the Kazusa database.

To facilitate simplicity in downstream PCR reactions of many selections in different viral strains, we incorporated a barcoded region directly after the stop codon, similar to a barcoding strategy used in H1N1 influenza viruses previously^48^. Here, however, we incorporated two barcodes separated by a cloning scar. Immediately after the stop codon are defined barcodes. The first four nucleotides of this 12-nucleotide barcode define a particular background virus (virus barcode), and the latter eight nucleotides define which position is mutated at that position (position barcode (**Figure S1B**). Both virus and position barcodes have a hamming distance of 3 or greater to prevent sequencing errors from misassigning barcodes, which was achieved using BARCOSEL^49^. These defined barcodes were added during PCR-based generation of the library (Twist Bioscience). During cloning (see below), a random 12-nucleotide barcode was added after a four base pair scar immediately after the defined barcodes. Following the random barcodes is a repeated 80 nt packing sequence as previously described^48,50^. The C-terminal region of the coding sequence was modified as previously described to prevent a repeat packing signal^48^, and the nucleotide sequence of the barcoded region was made consistent between all strains such that the primer binding sites enable amplification of the barcoding regions of all three strains (the assembled library sequences can be found at https://github.com/dpmaurer/flu_ab_escape). Linear double stranded DNA libraries with this design were ordered from Twist Bioscience.

### HA library cloning

We first generated a modified version of pHW2000^51^ where we replaced the stuffer sequence with a GFP cassette (pDM_RG068) (plasmid maps can be found at https://github.com/dpmaurer/flu_ab_escape). Bacterial colonies carrying these plasmids fluoresce green under blue light, allowing us to quantify the proportion of colonies carrying undigested plasmid after cloning. We used a three-piece golden gate cloning^52^ method to assemble the HA coding sequence library (Twist) and a DNA segment encoding the random 12-nucleotide barcode and repeated packing sequence (IDT) with the pDM_RG068 backbone. The linear DNA libraries were delivered in plates where each well contained variants at one given position. For each library, we resuspended each well in 5 µl of water and pooled 2 µl from all wells. Although in theory golden gate cloning does not require pre-digestion of the library or plasmid, we found cloning was most efficient when we digested the library and plasmid with BsmBI-v2 (NEB, #R0739L) (but not the barcode fragment due to the quantity) and purified these fragments by gel extraction prior to assembly. The three fragments were quantified by Qubit (ThermoFisher) and combined in a plasmid:library:barcode fragment ratio of 1:2:2 or 30:60:60 fmol. BsmBI-v2 golden gate assembly mix (NEB, #E1602S, 1.2µl) and the supplied T4 DNA ligase reaction buffer (1.2 µl) were added in a total reaction volume of 12 µl. The reaction was then cycled overnight between 42 °C (5 min) and 16 °C (5 min) a total of 60 times, followed by 60 °C (5 min), and 80 °C (20 min). The reaction was dialyzed by filling a Petrie dish with water, placing a piece of 0.025 µm filter paper on top, and pipetting the 12 µl reaction on top of the filter paper. The dialysis was allowed to proceed for 30 minutes at room temperature. The dialyzed sample was then removed and split equally in half to generate two replicate libraries. For each replicate library, we electroporated 100 µl of NEB 10-beta (NEB, #C3020K) in a BioRad electroporator using the preset E. coli protocol (2mm cuvette, 2.5 kV). After electroporation, E. coli transformations were recovered with 4 ml of the supplied media at 30 °C for 1 hour. The transformations were then diluted in a half-log series and the entire volume of each dilution was plated on 245 mm square agar plates (Corning, #431111) containing carbenicillin. In practice, we found it is only necessary to plate dilutions corresponding to 10^1.5^ and 10^2.0^, but also plating 10^2.5^ helps with the ease of counting colonies. The plates were allowed to dry before incubating at 30 °C for 24 hours. Then, colonies were counted and a goal of ∼100,000-200,000 colonies were scraped into 200 ml of LB with carbenicillin and grown for an additional 2 hours at 30 °C prior to maxi prepping the plasmid DNA (Macheregy-Nagel, #740414.50).

### Long read sequencing preparation (PacBio)

Maxi-prepped plasmids containing the replicate barcoded libraries were digested using SalI-HF (NEB) to linearize the plasmid prior to long-read sequencing. The linearized plasmids were prepped for sequencing by the Bauer Core using a SMRTbell v3.0 kit. The resulting libraries were sequenced using a PacBio Sequel IIe (Bauer Core). We implemented computational methods from previous reports of SARS-CoV-2 deep mutational scanning experiments^53,54^ to link barcode regions with individual variants.

### Virus library generation

Previous work has shown that reducing the bottleneck of an eight-plasmid reverse genetics system can increase the library generation efficiency^48,55^. To this end, we generated a 6-in-1 ∼18.1 kbp plasmid containing bidirectional cassettes of mouse-adapted A/Puerto Rico/8/1934 (PR8) PB2, PB1, PB1, PA, NP, M, NS. To generate this plasmid, we first removed the PaqCI site from pHW195 (PR8 NP, a gift from Jesse Bloom) by site-directed mutagenesis (Agilent). Then, we amplified the bidirectional cassettes with Q5 (NEB, #M0491L) and the pHW2000 plasmid backbone such that the whole plasmid could be assembled in a one-pot golden gate reaction with PaqCI (NEB, #R0745S). Due to repeated regions, we found it necessary to propagate this plasmid in NEB Stable E. coli (NEB, #C3040H) at 30 °C in LB medium, using non-baffled flasks. Maxi preps of this plasmid were always sequence confirmed by whole plasmid sequencing (Primordium Labs).

To avoid jackpotting or bottlenecking effects, we transfected each replicate of each library in 96-144 transfections in 6-well plates. A mixture of 600,000 HEK293T and 100,000 hCK cells per well were seeded in poly-D-lysine coated 6-well plates (Greiner, #657940). The next day when cells were ∼60-80% confluent, each well was transfected dropwise with the following after a 15 minute room temperature incubation: 200 µl Opti-MEM containing 500 ng HA library plasmid, 500 ng NA plasmid for the particular strain, 3 µg pDM_RG070 (6-in-1 PR8 plasmid), 500 ng hTMPRSS2 plasmid^56^, and 9 µl of TransIT-LT1 (MirusBio, #2306, 2:1 ratio). Transfections were performed in batches so that transfection mixtures were not sitting too long. After ∼16-18 hours, the FBS-containing media was removed, the cells were washed with PBS, and flu media containing 1 µg/ml TPCK-trypsin was added. Approximately 48 hours after replacing the media when cytopathic effect was apparent, the supernatants were pooled, clarified by centrifugation (800 xg, 5 min), aliquoted, and frozen at -80 °C to bank the “P0” stocks.

To enhance genotype-phenotype linkage, we passaged virus at a low MOI (0.01) across approximately 200-300 million cells. To do this, we seeded 4000 cm^2^, 16-layered flasks (Greiner, #678116) with ∼50,000 hCK cells per cm^2^ in flu media. The next day, we infected these cells with 3 million infectious particles in 500 mls of flu media. After 1 hour, we removed the inoculum, washed the cells with PBS, and replaced the media with fresh flu media containing 1 µg/ml TPCK-trypsin. ∼20-24 hours post-infection, we clarified the supernatant and aliquoted “P1” viral stocks. These P1 stocks were used for escape selections we describe here.

### Titering and neutralization assays

To titer or test neutralization against viral libraries, we used a standard ELISA-based microneutralization assay^57^ with some modifications. hCK cells were seeded in 96-well plates (Corning, #3595) at 20,000-30,000 cells per well in flu media. The next day, virus was prepared in half-log dilutions (for titering) or at 200 TCID50 / 50 µl (neutralization) in flu media containing 2 µg/ml TPCK-trypsin. Flu media (titering) or four-fold antibody dilutions (neutralization) were added to the viruses (1:1 ratio) and incubated at 37 °C / 5% CO2 for 1 hour. The media was then removed from the cells and virus (titering) or virus-antibody mixtures (neutralization) were overlayed on the hCK cells. After 1 hour at 37 °C / 5% CO2, the inoculum was removed, the cells were washed with PBS, and fresh flu media containing 1 µg/ml TPCK-trypsin was added. ∼18-20 hours later, the media was removed, cells were washed twice with PBS, and fixed with ice-cold 80% acetone in PBS for 10 minutes. After removing the fixative and air-drying, the plates were washed with wash buffer (PBS + 0.3% Tween20 (Sigma, #P1379)) three times before adding antibody diluent (PBS + 0.1% Tween20 + 1% BSA) to block. After blocking for 1 hour at room temperature, the plates were washed three times and a solution of anti-nucleoprotein antibodies (Sigma, #MAB8257 and #MAB8258, 1:4000 each) in antibody diluent were added for 1 hour at room temperature. After washing three times, a dilution of anti-mouse antibody HRP conjugate (seracare, #5220-0339, 1:20000) was added for 1 hour at room temperature. After washing five times, Ultra TMB was added (ThermoFisher, #34028) for ∼15 minutes). The development was stopped by addition of 2M sulfuric acid before reading at 450 nm (SpectraMax).

Titering calculations were performed by the Reed and Muench method^57,58^. Each dilution was performed in quadruplicate or eight replicates. Neutralization potencies were determined by subtracting background from cell only controls and normalizing 0% neutralization to the virus only controls. IC50 values were determined by a sigmoidal curve fit in GraphPad Prism. IC99 values were then determined by IC99 = (99/(100-99))^(1/hill^ ^slope)^ x IC50.

### Virus library selections

Escape selections were performed in T150 flasks after seeding 50,000 hCK cells per cm^2^ (∼7.5 million) the day prior. A number of infectious particles corresponding to an MOI of ∼0.05 were combined with antibody at 10x the IC99 value in a total of 28 mls of flu media containing trypsin. After incubation (37 °C / 5% CO2) for 1 hour, the hCK cells were washed with PBS, and new flu media containing trypsin was added. After ∼20-24 hours, supernatant was clarified by centrifugation and 20 mls were concentrated using 100K MWCO protein concentrators (ThermoFisher, #88532) by centrifugation (1000 xg, 4 °C, 30-60 min). The viral RNA from the supernatant was purified using QIAamp viral RNA mini preps (Qiagen, #52904). The final elution was 60 µl using the supplied buffer.

### Amplification of viral barcodes

To amplify the viral barcoded regions, potential DNA contamination was first removed by incubation of 8 µl viral RNA, 1 µl ezDNase, and 1 µl ezDNase buffer (ThermoFisher, #11766051) at 37 °C for 2 minutes. dNTPs (NEB, #N0447L, 1 µl), water (1 µl), and a primer that anneals in the 3’ noncoding region (prDM370, 2 µM, 1 µl, **Table S3**) were added and incubated at 65 °C for 5 minutes before placing on ice for 1 minute. SuperScript IV (ThermoFisher, #18090050, 1 µl), SuperScript IV buffer (4 µl), RNaseOUT (ThermoFisher, #10777019, 1 µl) and DTT (100 mM, 1 µl) were added to start cDNA synthesis by incubation at 50 °C for 30 minutes followed by 80 °C for 10 minutes. The cDNA was cleaned up by a 1.6X AMPure XP bead cleanup (Beckman, #A63881) with a final elution of 23 µl, of which 20 µl was recovered for the first PCR step.

To (1) increase the accuracy of amplicon sample assignment from NGS data, (2) increase the nucleotide diversity during sequencing, and (3) limit amplification bias, we amplified the barcoded region in a first PCR step with primers (**Table S3**) containing inline indices and cycled only four times^30^. This first PCR step was performed using a standard Q5 PCR: cDNA template (20 µl), Q5 buffer (10 µl), dNTPs (10 mM, 1 µl), water (13.5 µl), primers (10 mM, 2.5 µl each), and Q5 polymerase (0.5 µl) was combined. The reaction was cycled by 98 °C for 10 seconds; four cycles of 98 °C for 30s, 67 °C for 10s, 72 °C for 10s; 72 °C for 2 minutes. The reaction was purified with a 1.6X AMPure XP bead cleanup with a final elution of 23 µl, of which 20 µl was used for the second PCR step. To calculate functional scores^29^, we also amplified the barcoded region from the plasmid libraries using 10 ng as input for the first PCR step.

The second PCR step added NEBNext unique dual index primers onto the ends of the amplicon (NEB, #E6440S). This was achieved by a two-step PCR reaction containing the same reagents as in the first PCR step except the primers were 5 µl of the supplied unique dual index primer mix. The cycling conditions were the following: 98 °C for 30s; 14 cycles of 98 °C for 10s, 72 °C for 15s; 72 °C for 2 minutes. We found that increasing cycles above 14 often resulted in the formation of “bubble” products due to over amplification of the amplicon. The approximate concentrations were determined by D1000 Tape on a TapeStation (Agilent). The amplicons were pooled at equimolar concentrations and quantified using Qubit (ThermoFisher) prior to submission for Illumina sequencing using a NovaSeq SP flow cell (Bauer Core).

### ELISA

Recombinant full-length HA ectodomains or heads were coated on 96-well plates (Corning, #9018) in PBS at 2 µg/ml (full-length) or 5 µg/ml (head) (100 µl, overnight, 4 °C). The next day, wells were blocked with PBS containing 1% BSA and 0.1% Tween20 for 1 hour at room temperature. The blocking buffer was removed and antibodies (40 µl) were added for 1 hour at room temperature. The antibodies were removed and the plates were washed three times with PBS + 0.1% Tween20 before adding anti-human IgG HRP conjugate (Abcam, #ab97225, 1:20000) for 1 hour at room temperature. After washing three times with wash buffer, ABTS (ThermoFisher, #37615, 150 µl) was added. Absorbance was read at 405 nm (Molecular Devices, SpectraMax iD5). Wells that contained no primary antibody were used as a background measurement, which was subtracted from all values on the plate. KD values were determined by a sigmoidal fit in Prism 10 (GraphPad).

### Structure determination

UCA860 Fab and MA90 head were complexed at a Fab:HA head ratio of 4:1 and CH67 with MA90-G189E at a ratio of 2:1 at room temperature for 30 minutes. The complexes were concentrated to ∼12 mg/ml prior to crystallization. The complexes were crystallized by the hanging drop method in 96-well plates (Greiner, #655101) with ViewDrop II plate seals (sptlabtech, #4150-05600). The UCA860:MA90 and two CH67:MA90-G189E complexes crystallized in 20% PEG 4K / 200 mM imidazole / pH 7.4 / HCl, 10% PEG 8K / 100 mM HEPES / pH 7 / 500 mM NaCl, and 20% 8K / 100 mM HEPES / pH 7 / 100 mM MgSO4, respectively (**Table S1**). (+/-)-2-methyl-2,4-pentanediol (MPD) was added to mother liquor solutions at 12%, and 1 µl was used to harvest crystals before flash-cooling in liquid nitrogen.

Diffraction data was collected at Advanced Photon Source beam line 24-ID-E and processed using XDSGUI (https://strucbio.biologie.uni-konstanz.de/xdswiki/index.php/XDSGUI). Structures were solved by molecular replacement using PHASER^59^ in the PHENIX GUI^60^. The UCA860:MA90 structure was solved using the CH67 VHVL with the CDRH3 deleted and SI06 head models from PDB 4HKX^26^. The CH67:MA90-G189E structures were solved using the CH67 VHVL and CHCL from PDB 4HKX Structure refinement was done in PHENIX; coordinates and B factors were refined and the CDRH3 was manually built into density in COOT^61^. In the final round of refinement of the UCA860:MA90 structure, waters and translation libration screw (TLS) refinements were used. For the CH67-MA90-G189E structures, Ramachandran restraints were included during refinement in PHENIX. Notably, the HA head in the UCA860:MA90 complex had very large B factors for HA positions 54-59 and 71-90, which coincide with placement of N-linked glycans (N58 and N91). The MA90 head in the UCA860:MA90 complex was produced in HEK293F cells, such that full glycosylation is expected at N58 and N91. The CH67:MA90-G189E (1) complex showed clear density for the HA head and variable antibody regions, but poor density for the constant domains, although they were correctly placed during molecular replacement. Due to the poor constant region density, the overall crystallographic statistics are relatively poor for a structure at this resolution despite clear density for the HA head and variable domains (**Table S1**). The structures of UCA860:MA90 and CH67:MA90-G189E complexes have been uploaded to the PDB as 9AZR, 9AZT, and 9AZV.

### Computational methods

The computational methods used here are largely based on previous computational analyses from deep mutational scanning experiments^9,29,53^. These methods detail barcode linking with PacBio sequencing data^53^, functional score calculations^29^, and differential selection calculations^9^. We also used packages *alignparse*^62^ and *dms_variants* (https://jbloomlab.github.io/dms_variants/). Jupyter notebooks and outputs for the code described below are available at https://github.com/dpmaurer/flu_ab_escape.

#### Linking barcodes to variants

The PacBio data generated as above was used to link the barcodes with respective variants. We used *alignparse*^62^ version 0.6.2, based on previously written code from Starr et al.^53^, to link the barcoded region to individual variants. We removed variants with insertions but not deletions as we intended to include deletions in our libraries. For each library, >10X coverage of the codon variant library size had at least 3 PacBio CCSs per barcode.

#### Barcode count analysis

We merged paired end reads using PEAR^63^. From merged reads, we filtered for the expected length based on the amplicon, inline indices, and UMIs. We additionally filtered out sequences containing ambiguous base calls or those that did not match the expected inline indices. From these filtered reads, we combined UMIs from each side of the amplicon into a single UMI and counted unique barcode-UMI reads.

#### Functional score calculation

We determined functional scores using *dms_variants* (https://jbloomlab.github.io/dms_variants/), which calculates the base 2 log enrichment before and after selection compared to wild-type. In our case, pre-selection was the plasmid library and post-selection was after generation and a low MOI passage. We calculated functional score by amino acid substitutions and calculated synonymous variants separately from wild-type.

#### Differential selection calculation

We determined escape by calculating differential selection^9^ (https://jbloomlab.github.io/dms_tools2/diffsel.html). This metric calculates the enrichment of each amino acid variant compared to wild-type between a selection and a “mock” selection (here, a passage without added antibody). A pseudocount (we used a pseudocount of 10) is added to each variant count that is scaled by read depth for a certain position. For analyses, we only considered those variants that were positively selected (differential selection > 0) in both biological replicates and all other values were set to 0. For each selection, we normalized the maximum differential selection value to 1. We defined escape mutations as those that were >0.5 in both replicates. After normalization, we summed the differential selection values at each site (**Figure S1D**).

### Data and code availability

Raw PacBio and Illumina sequencing data have been deposited to NCBI under BioProject PRJNA1056469. All code used to analyze the sequencing data and the resulting output files used to generate the figures at https://github.com/dpmaurer/flu_ab_escape. Structures have been deposited to the PDB under accession codes 9AZR, 9AZT, and 9AZV.

