## Supplementary Materials for "Antigenic drift expands viral escape pathways from imprinted host humoral immunity"

**Table S1. X-ray crystallography data collection and refinement statistics.**

| | **UCA860 Fab bound to H1 A/Massachusetts/1/1990 head** | **CH67 Fab bound to H1 A/Massachusetts/1/1990 head G189E mutant (1)** | **CH67 Fab bound to H1 A/Massachusetts/1/1990 head G189E mutant (2)** |
|---|---|---|---|
| **Data Collection** | | | |
| Resolution (Å) | 48.94 - 2.30 (2.44 - 2.30) | 47.2 - 2.94 (3.12-2.94) | 47.03 - 3.16 (3.35 - 3.16) |
| Wavelength (Å) | 0.97918 | 0.97918 | 0.97918 |
| Space Group | C121 | $P2_12_12$ | $P2_12_12$ |
| Unit cell dimensions (a, b, c) (Å) | 202.543, 71.863, 65.402 | 80.75, 91.7, 110.1 | 76.28, 93.47, 108.85 |
| Unit cell angles (α, β, γ) (°) | 90, 104.867, 90 | 90, 90, 90 | 90, 90, 90 |
| I/σ | 10.35 (2.15) | 22.92 (2.15) | 10.81 (2.18) |
| $R_{meas}$ (%) | 13.8 (93.0) | 6.9 (98.9) | 14.8 (81.0) |
| $CC_{1/2}$ (%) | 99.5 (91.4) | 100 (93.2) | 99.6 (90.1) |
| Completeness (%) | 98.0 (90.8) | 99.2 (95.1) | 99.7 (99.0) |
| Number of observed reflections | 266351 (42654) | 230525 (35152) | 80958 (12964) |
| Number of unique reflectiions | 39860 (5923) | 17764 (2695) | 13883 (2181) |
| Redundancy | 6.7 (7.20) | 6.6 (6.6) | 5.8 (5.9) |
| **Refinement** | | | |
| Resolution (Å) | 48.31 - 2.301 (2.383 - 2.301) | 47.2 - 2.945 (3.05 - 2.945) | 46.73 - 3.16 (3.273 - 3.16) |
| Reflections used in refinement | 39803 (3535) | 17746 (1629) | 13830 (1350) |
| Reflections used for $R_{free}$ | 2006 (192) | 1774 (163) | 1382 (133) |
| $R_{work}$ (%) | 22.87 (31.99) | 29.41 (48.15) | 23.69 (34.83) |
| $R_{free}$ (%) | 27.03 (35.27) | 33.76 (49.04) | 28.71 (43.11) |
| Ramachandran favored/allowed (%) | 95.82 / 4.18 | 94.77 / 4.75 | 93.94 / 5.59 |
| Ramachandran outliers (%) | 0 | 0.48 | 0.47 |
| Rmsd bond lengths (Å) | 0.003 | 0.004 | 0.002 |
| Rmsd bond angles (°) | 0.56 | 0.76 | 0.49 |
| Average B-factor | 54.46 | 139.78 | 96.79 |
| **Crystallization Conditions** | 20% PEG 4k, 200 mM imidazole, pH 7.4, HCl | 10% PEG 8K, 100 mM HEPES, pH 7, 500 mM NaCl | 20% PEG 8K, 100 mM HEPES, pH 7, 100 mM MgSO4 |
| **PDB ID** | 9AZR | 9AZT | 9AZV |

**Table S2. Selected codon mutations for each amino acid in the viral libraries.**

| Amino Acid | Codon |
|---|---|
| A | GCA |
| C | TGC |
| D | GAT |
| E | GAA |
| F | TTC |
| G | GGA |
| H | CAT |
| I | ATA |
| K | AAA |
| L | CTT |
| M | ATG |
| N | AAT |
| P | CCA |
| Q | CAA |
| R | AGA |
| S | TCA |
| T | ACA |
| V | GTG |
| W | TGG |
| Y | TAT |
| * | TAA |

### Table S3. Primers for amplification of barcoded viral RNA.

| Primer Name | Primer Description | Sequence (5' to 3') | Inline index |
|---|---|---|---|
| prDM370 | RT primer | AGCAAAAGCAGGGGAAAATAAAAACAACC | N/A |
| prDM382 | Fwd primer for PCR1 with inline index and UMI | ACACTCTTTCCCTACACGACGCTCTTCCGATCT(N:25252525)(N)(N)(N)(N)(N)(N)(N)TTGATCGAGCCTCCAGTGTAGGATTTGC | TTGATCG |
| prDM383 | Fwd primer for PCR1 with inline index and UMI | ACACTCTTTCCCTACACGACGCTCTTCCGATCT(N:25252525)(N)(N)(N)(N)(N)(N)(N)CATTCTCGAGAGCCTCCAGTGTAGGATTTGC | CATTCTCGAG |
| prDM384 | Fwd primer for PCR1 with inline index and UMI | ACACTCTTTCCCTACACGACGCTCTTCCGATCT(N:25252525)(N)(N)(N)(N)(N)(N)(N)CCTTAGTACTAAGCCTCCAGTGTAGGATTTGC | CCTTAGTACTA |
| prDM385 | Fwd primer for PCR1 with inline index and UMI | ACACTCTTTCCCTACACGACGCTCTTCCGATCT(N:25252525)(N)(N)(N)(N)(N)(N)(N)AGTGGAATACCTAGCCTCCAGTGTAGGATTTGC | AGTGGAATACCT |
| prDM386 | Fwd primer for PCR1 with inline index and UMI | ACACTCTTTCCCTACACGACGCTCTTCCGATCT(N:25252525)(N)(N)(N)(N)(N)(N)(N)GGTATCAAGCCTCCAGTGTAGGATTTGC | GGTATCA |
| prDM387 | Fwd primer for PCR1 with inline index and UMI | ACACTCTTTCCCTACACGACGCTCTTCCGATCT(N:25252525)(N)(N)(N)(N)(N)(N)(N)GATGTCCTAGCCTCCAGTGTAGGATTTGC | GATGTCCT |
| prDM388 | Fwd primer for PCR1 with inline index and UMI | ACACTCTTTCCCTACACGACGCTCTTCCGATCT(N:25252525)(N)(N)(N)(N)(N)(N)(N)CTCATCCAGAGCCTCCAGTGTAGGATTTGC | CTCATCCAG |
| prDM389 | Fwd primer for PCR1 with inline index and UMI | ACACTCTTTCCCTACACGACGCTCTTCCGATCT(N:25252525)(N)(N)(N)(N)(N)(N)(N)GACGGAACTCAGCCTCCAGTGTAGGATTTGC | GACGGAACTC |
| prDM390 | Fwd primer for PCR1 with inline index and UMI | ACACTCTTTCCCTACACGACGCTCTTCCGATCT(N:25252525)(N)(N)(N)(N)(N)(N)(N)ATCGCAGGCATAGCCTCCAGTGTAGGATTTGC | ATCGCAGGCAT |
| prDM391 | Fwd primer for PCR1 with inline index and UMI | ACACTCTTTCCCTACACGACGCTCTTCCGATCT(N:25252525)(N)(N)(N)(N)(N)(N)(N)TCACGACTAGTAAGCCTCCAGTGTAGGATTTGC | TCACGACTAGTA |
| prDM392 | Rev primer for PCR1 with inline index and UMI | GACTGGAGTTCAGACGTGTGCTCTTCCGATCT(N:25252525)(N)(N)(N)(N)(N)(N)(N)TACCTGACCCCAGGGAGACCAAAAG | TACCTGA |
| prDM393 | Rev primer for PCR1 with inline index and UMI | GACTGGAGTTCAGACGTGTGCTCTTCCGATCT(N:25252525)(N)(N)(N)(N)(N)(N)(N)AGAACCATCCCCAGGGAGACCAAAAG | AGAACCAT |
| prDM394 | Rev primer for PCR1 with inline index and UMI | GACTGGAGTTCAGACGTGTGCTCTTCCGATCT(N:25252525)(N)(N)(N)(N)(N)(N)(N)ACCGTTGACCCCCAGGGAGACCAAAAG | ACCGTTGAC |
| prDM395 | Rev primer for PCR1 with inline index and UMI | GACTGGAGTTCAGACGTGTGCTCTTCCGATCT(N:25252525)(N)(N)(N)(N)(N)(N)(N)GCAAGCTGTTCCCCAGGGAGACCAAAAG | GCAAGCTGTT |
| prDM396 | Rev primer for PCR1 with inline index and UMI | GACTGGAGTTCAGACGTGTGCTCTTCCGATCT(N:25252525)(N)(N)(N)(N)(N)(N)(N)TCGGTGGTACGCCCCAGGGAGACCAAAAG | TCGGTGGTACG |
| prDM397 | Rev primer for PCR1 with inline index and UMI | GACTGGAGTTCAGACGTGTGCTCTTCCGATCT(N:25252525)(N)(N)(N)(N)(N)(N)(N)GTTCAACCGATTCCCCAGGGAGACCAAAAG | GTTCAACCGATT |
| prDM398 | Rev primer for PCR1 with inline index and UMI | GACTGGAGTTCAGACGTGTGCTCTTCCGATCT(N:25252525)(N)(N)(N)(N)(N)(N)(N)CGTACAACCCCAGGGAGACCAAAAG | CGTACAA |
| prDM399 | Rev primer for PCR1 with inline index and UMI | GACTGGAGTTCAGACGTGTGCTCTTCCGATCT(N:25252525)(N)(N)(N)(N)(N)(N)(N)GAGATCACCCCCAGGGAGACCAAAAG | GAGATCAC |
| NEBNext i5 | Fwd primer for PCR2 from NEBNext #6440, X denotes index | AATGATACGGCGACCACCGAGATCTACACXXXXXXXXACACTCTTTCCCTACACGACGCTCTTCCGATCT | N/A |
| NEBNext i7 | Rev primer for PCR2 from NEBNext #6640, X denotes index | CAAGCAGAAGACGGCATACGAGATXXXXXXXXGTGACTGGAGTTCAGACGTGTGCTCTTCCGATCT | N/A |

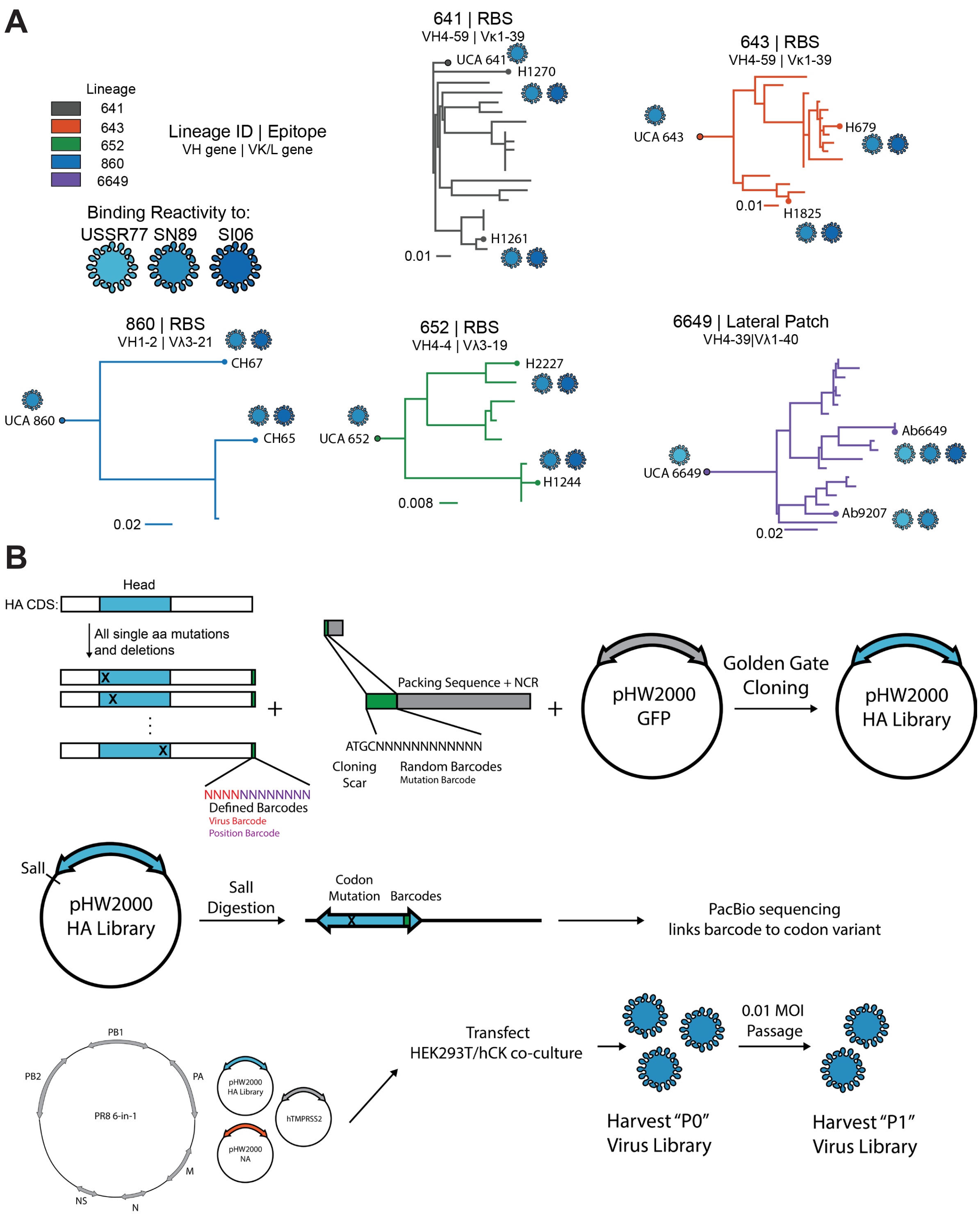
A
Lineage
641
643
652
860
6649
Lineage ID | Epitope
VH gene | VK/L gene
Binding Reactivity to:
USSR77 SN89 SI06
641 | RBS
VH4-59 | Vκ1-39
UCA 641
H1270
H1261
0.01
643 | RBS
VH4-59 | Vκ1-39
UCA 643
H679
0.01
H1825
860 | RBS
VH1-2 | Vλ3-21
CH67
UCA 860
CH65
0.02
652 | RBS
VH4-4 | Vλ3-19
H2227
UCA 652
H1244
0.008
6649 | Lateral Patch
VH4-39|Vλ1-40
Ab6649
UCA 6649
Ab9207
0.02
B
Head
HA CDS:
All single aa mutations and deletions
NNNNNNNNNNNNN
Defined Barcodes
Virus Barcode
Position Barcode
Packing Sequence + NCR
ATGCNNNNNNNNNNNN
Cloning Scar
Random Barcodes
Mutation Barcode
pHW2000 GFP
Golden Gate Cloning
pHW2000 HA Library
SalI
pHW2000 HA Library
SalI Digestion
Codon Mutation
Barcodes
PacBio sequencing links barcode to codon variant
PB1
PB2
PA
PR8 6-in-1
M
NS
N
pHW2000 HA Library
hTMPRSS2
pHW2000 NA
Transfect HEK293T/hCK co-culture
0.01 MOI Passage
Harvest "P0" Virus Library
Harvest "P1" Virus Library

**C**

```
        52                  62                  72                  82                  92                  102               111
        |                   |                   |                   |                   |                   |                 |
USSR77  A P L Q L G K C N I A G W I L G N P E C E S L F S K K S W S Y I A E T P N S E N G T C Y P G Y F A D Y E E L R E Q L S S
SN89    A P L Q L G N C S I A G W I L G N P E C E S L F S K E S W S Y I A E T P N S E N G T C Y P G Y F A D Y E E L R E Q L S S
SI06    A P L Q L G N C S V A G W I L G N P E C E L L I S R E S W S Y I V E K P N P E N G T C Y P G H F A D Y E E L R E Q L S S

        112                 122                 132                 141                 151                 161               170
        |                   |                   |                   |                   |                   |                 |
USSR77  V S S F E R F E I F P K E R S W P K H N V T R G V T A S C S H K G K S S F Y R N L L W L T E K N G S Y P N L S K S Y V N
SN89    V S S F E R F E I F P K E S S W P N H T V T K G V T A A C S H N G K S S F Y R N L L W L T E K N G L Y P N L S K S Y V N
SI06    V S S F E R F E I F P K E S S W P N H T T T - G V S A S C S H N G E S S F Y K N L L W L T G K N G L Y P N L S K S Y A N
                                                  |                       |                           |             |
                                                  133a                    145                         159           166

        171                 181                 191                 201                 211                 221               230
        |                   |                   |                   |                   |                   |                 |
USSR77  N K E K E V L V L W G V H H P S N I E D Q K T I Y R K E N A Y V S V V S S N Y N R R F T P E I A E R P K V R G Q A G R I
SN89    N K E K E V L V L W G V H H P S N I G D Q R A I Y H T E N A Y V S V V S S H Y S R R F T P E I A K R P K V R G Q E G R I
SI06    N K E K E V L V L W G V H H P P N I G D Q R A L Y H T E N A Y V S V V S S H Y S R K F T P E I A K R P K V R D Q E G R I
                                          |       |                                                                 |
                                          189     193                                                               226

        231                 241                 251                 261
        |                   |                   |                   |
USSR77  N Y Y W T L L E P G D T I I F E A N G N L I A P W H A F A L N R G
SN89    N Y Y W T L L E P G D T I I F E A N G N L I A P W Y A F A L S R G
SI06    N Y Y W T L L E P G D T I I F E A N G N L I A P R Y A F A L S R G
```

D

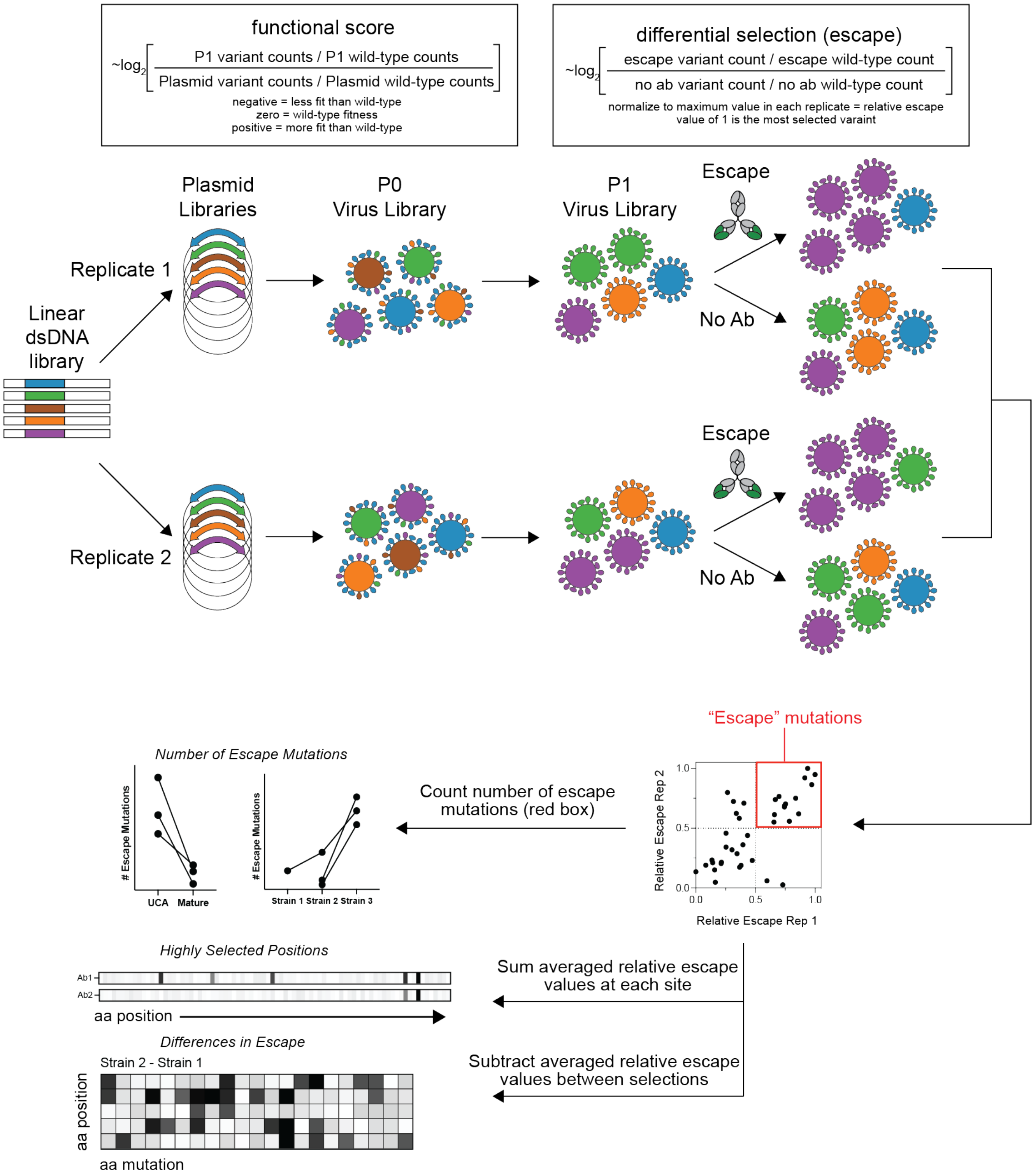

functional score
~log2 [ P1 variant counts / P1 wild-type counts / Plasmid variant counts / Plasmid wild-type counts ]
negative = less fit than wild-type
zero = wild-type fitness
positive = more fit than wild-type
differential selection (escape)
~log2 [ escape variant count / escape wild-type count / no ab variant count / no ab wild-type count ]
normalize to maximum value in each replicate = relative escape
value of 1 is the most selected varaint
Plasmid Libraries
P0 Virus Library
P1 Virus Library
Escape
No Ab
Replicate 1
Replicate 2
Linear dsDNA library
"Escape" mutations
Relative Escape Rep 2
Relative Escape Rep 1
Count number of escape mutations (red box)
Number of Escape Mutations
### Escape Mutations
UCA
Mature
Strain 1
Strain 2
Strain 3
Sum averaged relative escape values at each site
Highly Selected Positions
Ab1
Ab2
aa position
Subtract averaged relative escape values between selections
Differences in Escape
Strain 2 - Strain 1
aa position
aa mutation

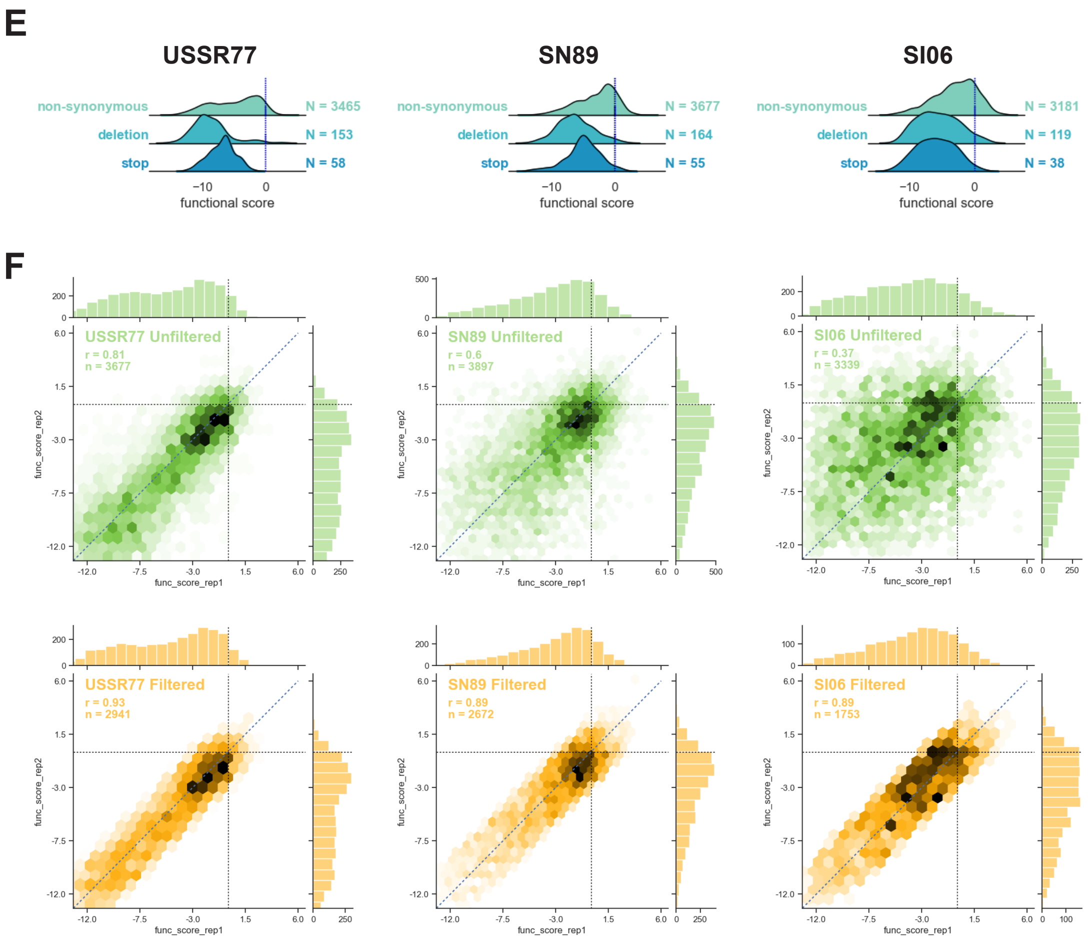


**Figure S1. Antibody lineage reconstructions and generation of virus libraries. (A)** Lineage reconstructions of four RBS lineages and one lateral patch lineage[22-26]. **(B)** Schematic describing the procedure to generate virus libraries: (1) all single amino acid mutations and deletions were introduced into the HA head region with defined barcodes appended at for each position, (2) golden gate cloning was used to append a random barcode along with cloning into a pHW2000-based[51] reverse genetics plasmid, (3) the plasmid library was digested with SalI prior to PacBio sequencing, (4) a plasmid carrying all six PR8 internal segments was co-transfected with the HA plasmid library, NA plasmid, and human TMPRSS2 plasmid into a co-culture of HEK293T/hCK cells and subsequently passaged at low MOI on hCK cells. **(C)** Amino acid alignment of the HA head regions (H3 numbering). **(D)** Overview of experimental design: (1) independent replicate DNA libraries were generated, (2) independent virus libraries were generated, (3) a low MOI passage supports genotype-phenotype linking, (4) escape is determined by passaging in the presence or absence of antibody, (5) computational analyses determine the functional score, number of escape mutations, highly selected positions, and differences in escape as outlined. **(D)** Functional scores of the "P1" viral libraries showing depletion of deletion and stop codons relative to non-synonymous mutations. Black and blue dotted lines represent wild-type and synonymous mutations, respectively, and are overlapping. **(E)** Correlations of functional scores from two biological replicate virus libraries for each strain. Pearson's correlation coefficient is shown. "n" is the number of amino acid mutations represented. Data filtered for a functional score difference of 3 between replicates.

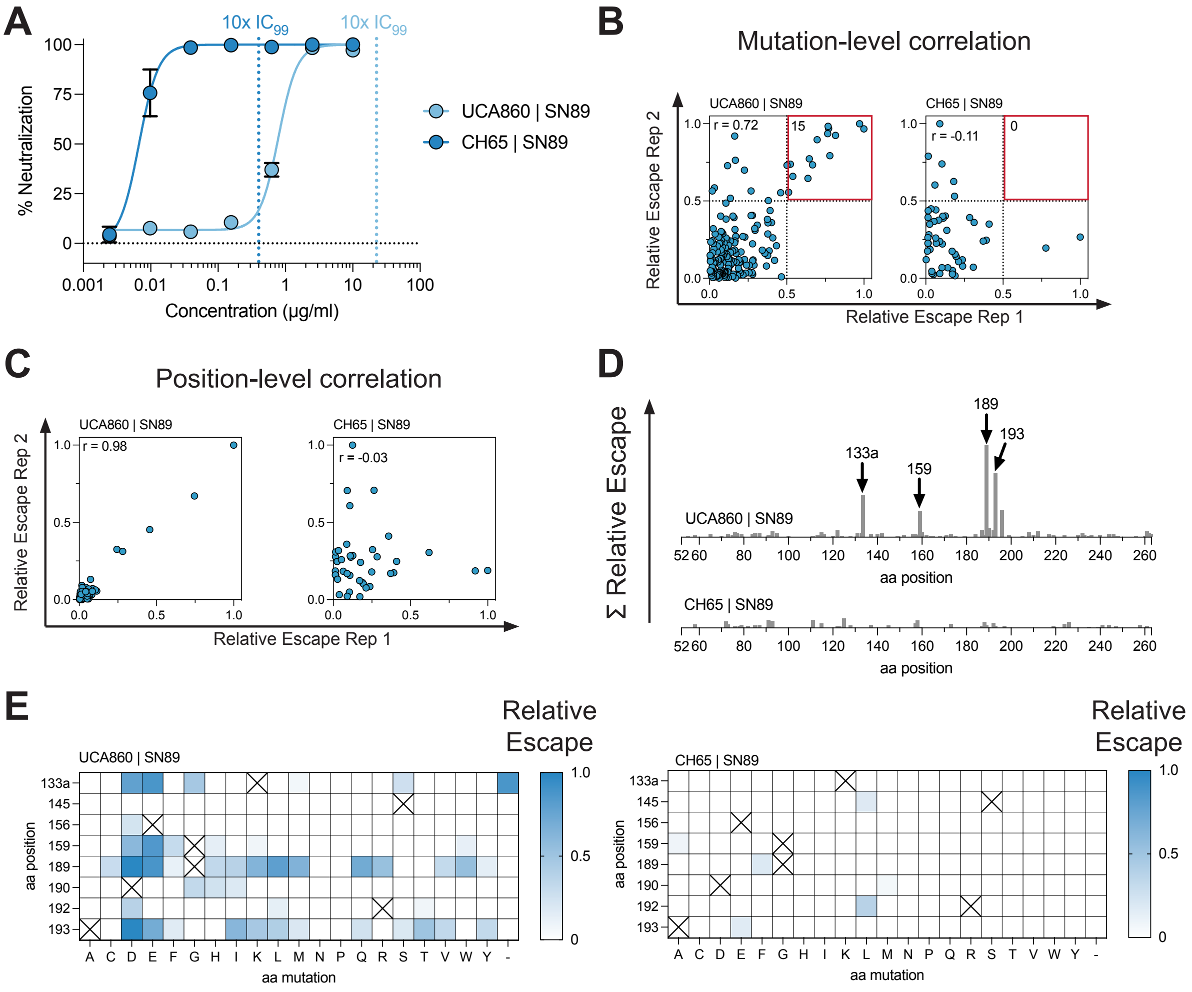
A
10x IC99
10x IC99
% Neutralization
Concentration (μg/ml)
UCA860 | SN89
CH65 | SN89
B
Mutation-level correlation
UCA860 | SN89
CH65 | SN89
r = 0.72
r = -0.11
15
0
Relative Escape Rep 2
Relative Escape Rep 1
C
Position-level correlation
UCA860 | SN89
CH65 | SN89
r = 0.98
r = -0.03
Relative Escape Rep 2
Relative Escape Rep 1
D
Σ Relative Escape
133a
159
189
193
UCA860 | SN89
CH65 | SN89
aa position
E
UCA860 | SN89
CH65 | SN89
Relative Escape
aa position
aa mutation

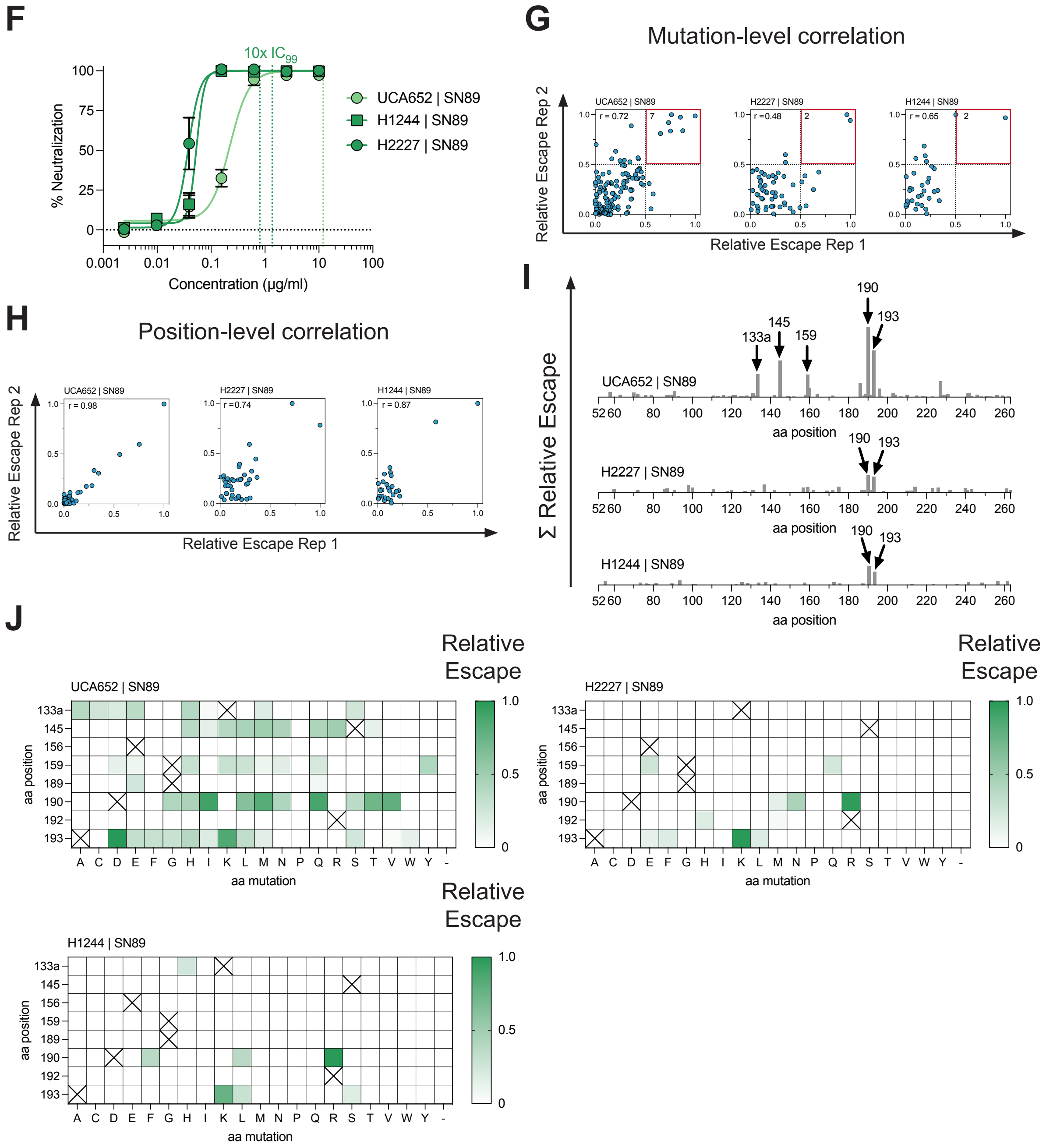
F
10x IC99
% Neutralization
Concentration (μg/ml)
UCA652 | SN89
H1244 | SN89
H2227 | SN89
G
Mutation-level correlation
Relative Escape Rep 2
Relative Escape Rep 1
H
Position-level correlation
I
Σ Relative Escape
aa position
J
Relative Escape
aa position
aa mutation

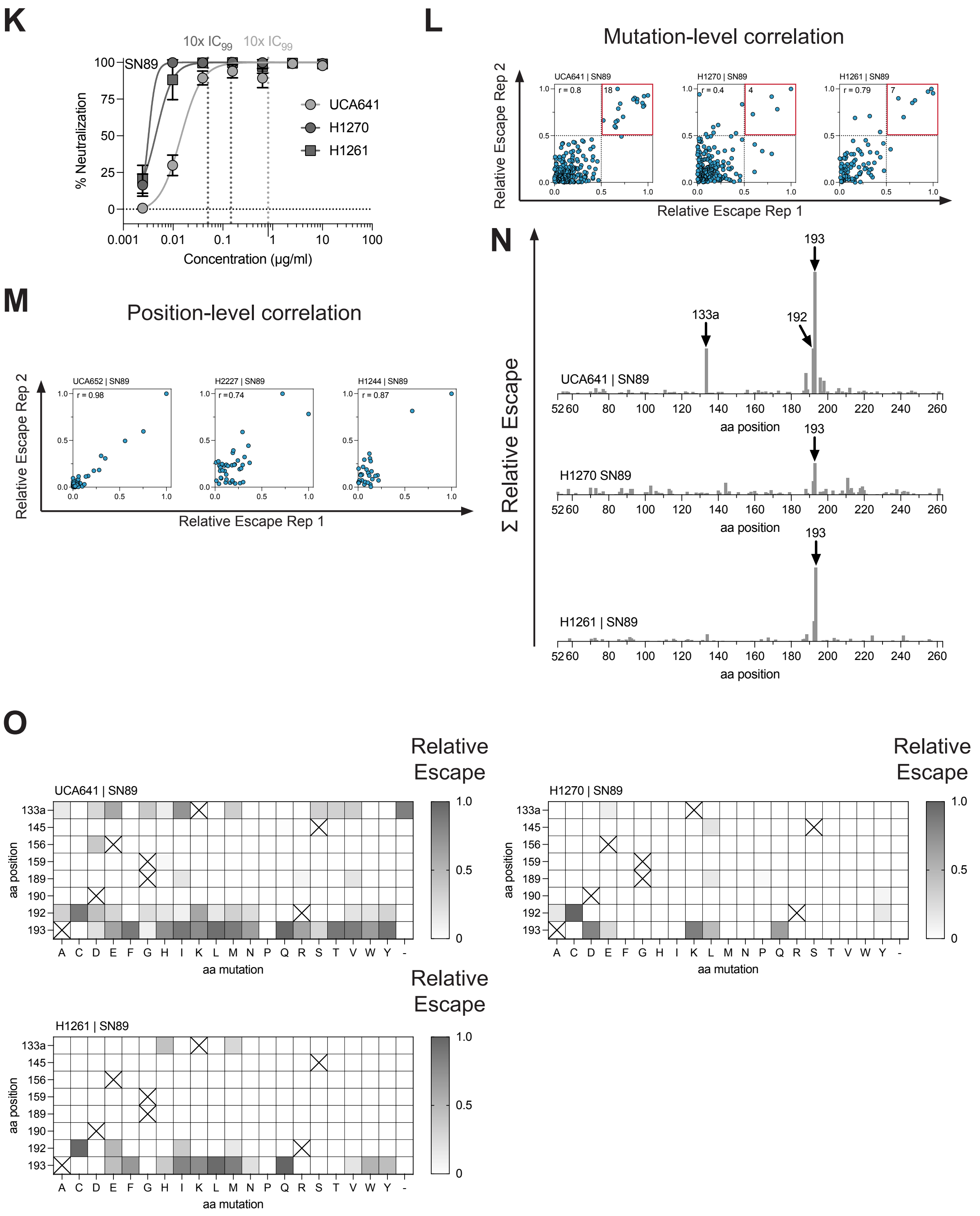

K
10x IC99
10x IC99
SN89
% Neutralization
Concentration (µg/ml)
UCA641
H1270
H1261
L
Mutation-level correlation
UCA641 | SN89
H1270 | SN89
H1261 | SN89
r = 0.8
r = 0.4
r = 0.79
Relative Escape Rep 2
Relative Escape Rep 1
M
Position-level correlation
UCA652 | SN89
H2227 | SN89
H1244 | SN89
r = 0.98
r = 0.74
r = 0.87
Relative Escape Rep 2
Relative Escape Rep 1
N
Σ Relative Escape
UCA641 | SN89
H1270 SN89
H1261 | SN89
133a
192
193
aa position
O
Relative Escape
UCA641 | SN89
H1270 | SN89
H1261 | SN89
aa position
aa mutation

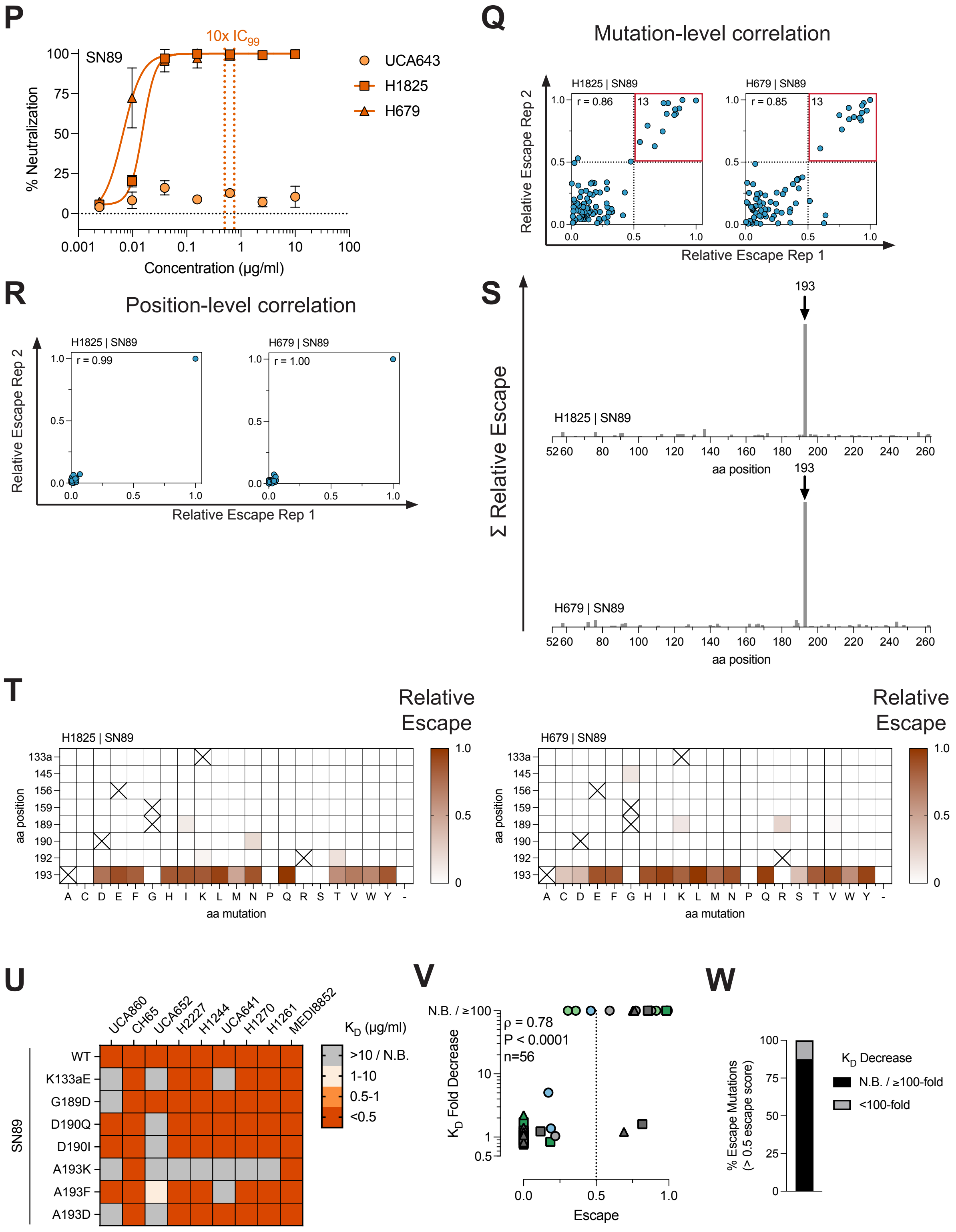


**Figure S2. Escape selections with RBS antibodies against the imprinting strain. (A,F,K,P)** Neutralization potencies for lineages 860, 652, 641, and 643. Dotted lines indicate the concentrations used in selections. **(B,G,L,Q)** Correlation between biological replicates for each mutation that shows positive differential selection in both replicates. Pearson's correlation coefficient is shown. Red box indicates our definition of escape mutations. The number inside the red box indicates the number of escape mutations. **(C,H,M,R)** Correlation between biological replicates summed by position. Pearson's correlation coefficient is shown. **(D,N,I,S)** Summed escape by position for all positions included in the libraries. All plots have a y-axis of 15. **(E,J,O,T)** Heatmaps showing the relative escape for each individual selection, highlighting the positions that were selected by RBS antibodies. **(U)** Avidities determined by ELISA for antibodies against SN89 variants. **(V)** Correlation of avidities shown in U with escape scores. **(W)** Proportion of mutations with escape scores >0.5 (our definition of escape mutation) that have over >100-fold $K_D$ decrease from SN89 escape selections.

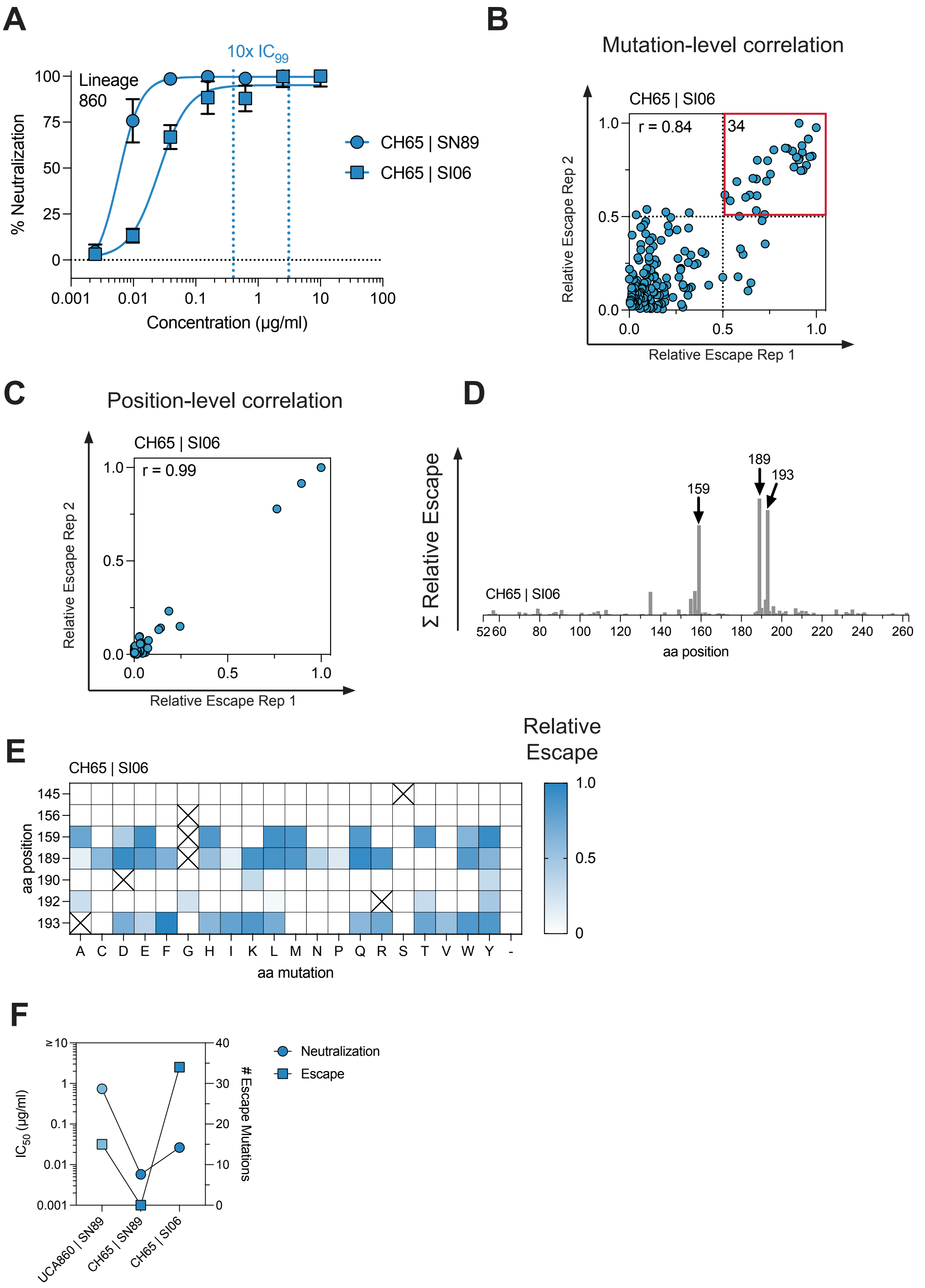
A
10x $IC_{99}$
Lineage 860
% Neutralization
100
75
50
25
0
0.001
0.01
0.1
1
10
100
Concentration (μg/ml)
CH65 | SN89
CH65 | SI06
B
Mutation-level correlation
CH65 | SI06
r = 0.84
34
Relative Escape Rep 2
Relative Escape Rep 1
0.0
0.5
1.0
C
Position-level correlation
CH65 | SI06
r = 0.99
Relative Escape Rep 2
Relative Escape Rep 1
0.0
0.5
1.0
D
Σ Relative Escape
CH65 | SI06
159
189
193
52
60
80
100
120
140
160
180
200
220
240
260
aa position
E
CH65 | SI06
Relative Escape
1.0
0.5
0
aa position
145
156
159
189
190
192
193
A C D E F G H I K L M N P Q R S T V W Y -
aa mutation
F
Neutralization
Escape
$IC_{50}$ (μg/ml)
≥10
1
0.1
0.01
0.001
### Escape Mutations
40
30
20
10
0
UCA860 | SN89
CH65 | SN89
CH65 | SI06

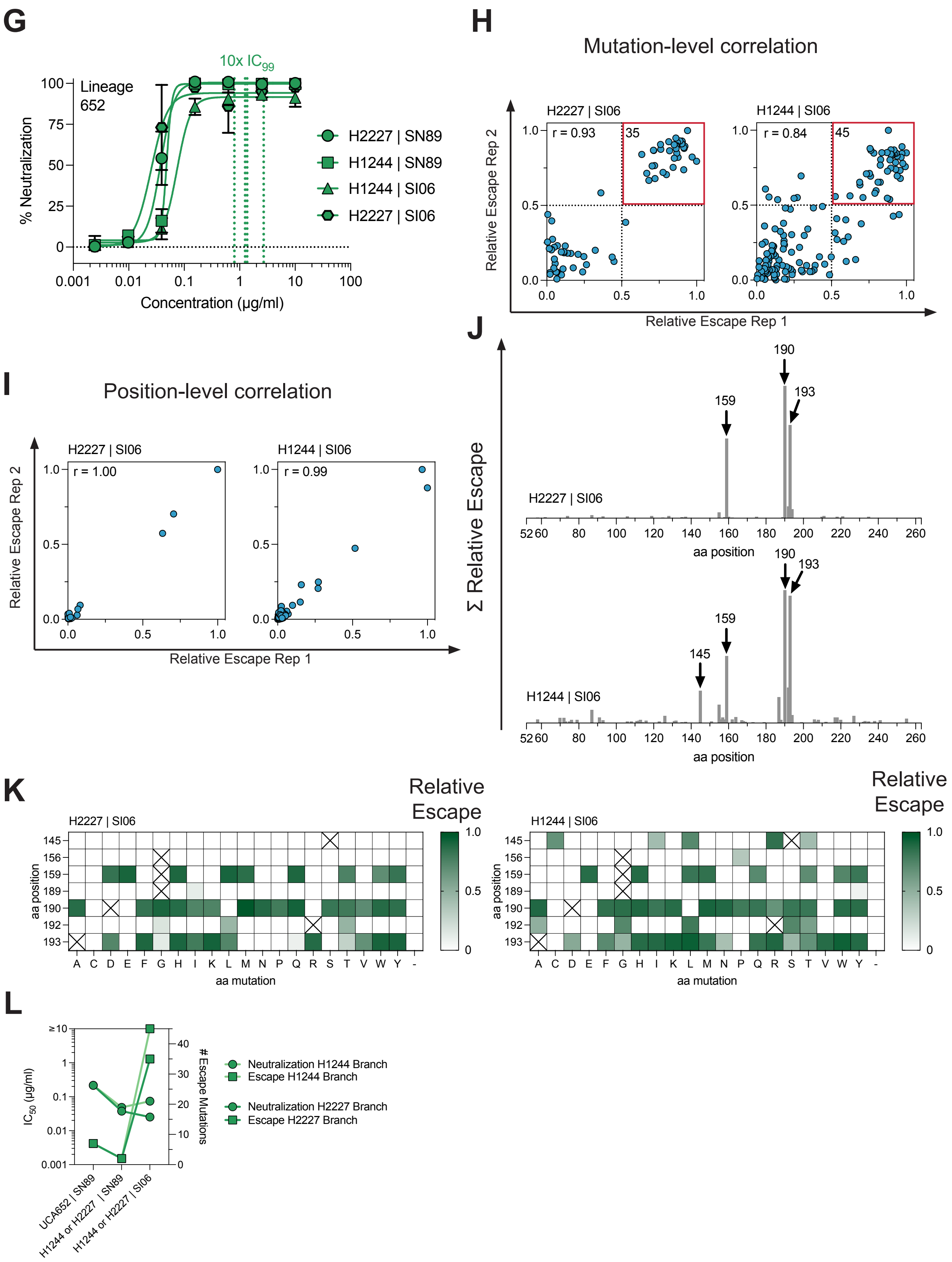
G
10x IC99
Lineage 652
% Neutralization
Concentration (µg/ml)
H2227 | SN89
H1244 | SN89
H1244 | SI06
H2227 | SI06
H
Mutation-level correlation
H2227 | SI06
H1244 | SI06
r = 0.93
r = 0.84
35
45
Relative Escape Rep 2
Relative Escape Rep 1
I
Position-level correlation
H2227 | SI06
H1244 | SI06
r = 1.00
r = 0.99
Relative Escape Rep 2
Relative Escape Rep 1
J
Σ Relative Escape
190
193
159
145
H2227 | SI06
H1244 | SI06
aa position
K
H2227 | SI06
H1244 | SI06
Relative Escape
aa position
aa mutation
L
IC50 (µg/ml)
### Escape Mutations
Neutralization H1244 Branch
Escape H1244 Branch
Neutralization H2227 Branch
Escape H2227 Branch
UCA652 | SN89
H1244 or H2227 | SN89
H1244 or H2227 | SI06

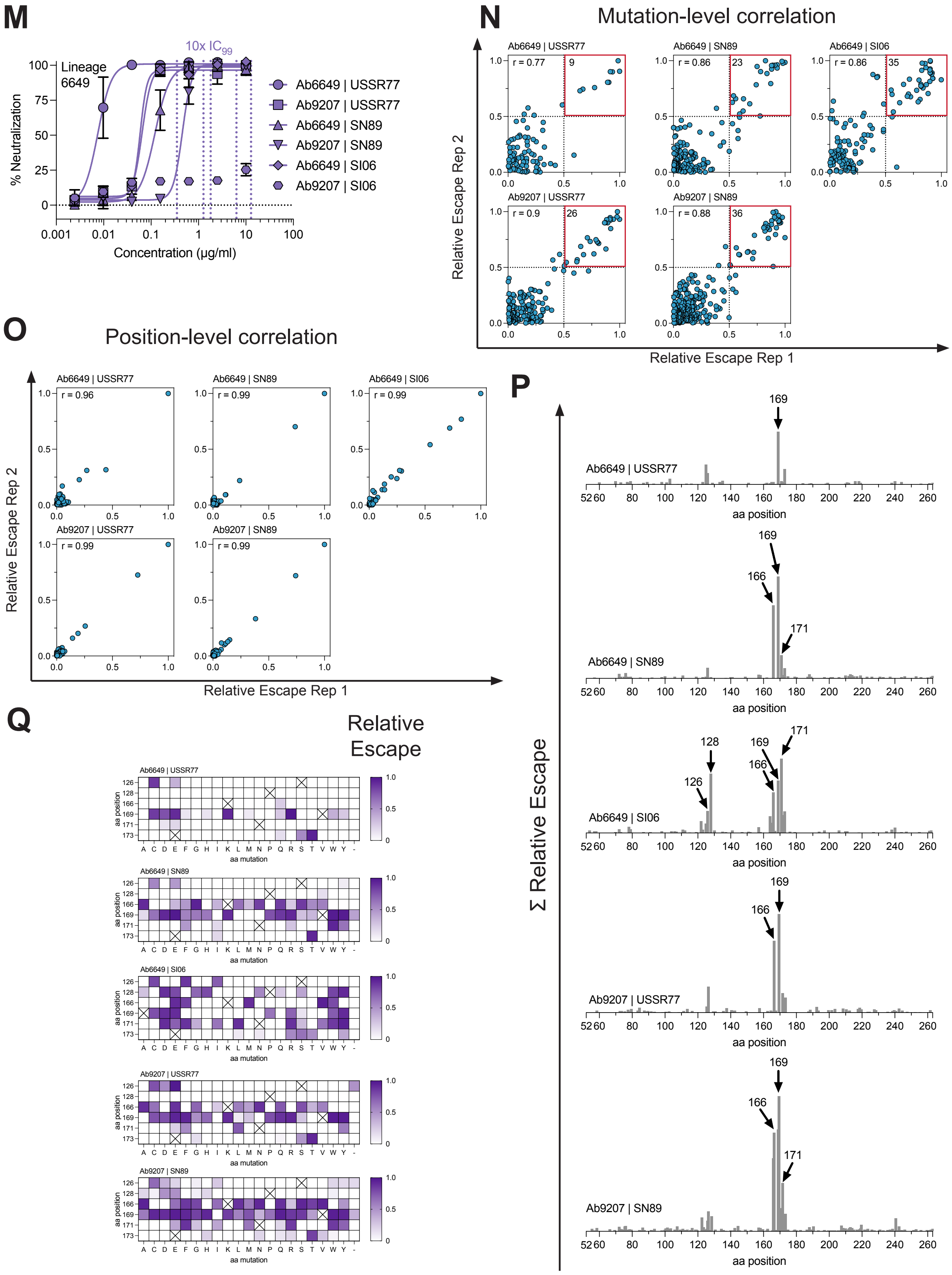
M
10x IC99
Lineage 6649
% Neutralization
Concentration (μg/ml)
Ab6649 | USSR77
Ab9207 | USSR77
Ab6649 | SN89
Ab9207 | SN89
Ab6649 | SI06
Ab9207 | SI06
N
Mutation-level correlation
Ab6649 | USSR77
r = 0.77
9
Ab6649 | SN89
r = 0.86
23
Ab6649 | SI06
r = 0.86
35
Ab9207 | USSR77
r = 0.9
26
Ab9207 | SN89
r = 0.88
36
Relative Escape Rep 2
Relative Escape Rep 1
O
Position-level correlation
Ab6649 | USSR77
r = 0.96
Ab6649 | SN89
r = 0.99
Ab6649 | SI06
r = 0.99
Ab9207 | USSR77
r = 0.99
Ab9207 | SN89
r = 0.99
Relative Escape Rep 2
Relative Escape Rep 1
P
Σ Relative Escape
Ab6649 | USSR77
169
Ab6649 | SN89
169
166
171
Ab6649 | SI06
128
126
169
166
171
Ab9207 | USSR77
169
166
Ab9207 | SN89
169
166
171
aa position
Q
Relative Escape
Ab6649 | USSR77
Ab6649 | SN89
Ab6649 | SI06
Ab9207 | USSR77
Ab9207 | SN89
aa position
aa mutation

**R**

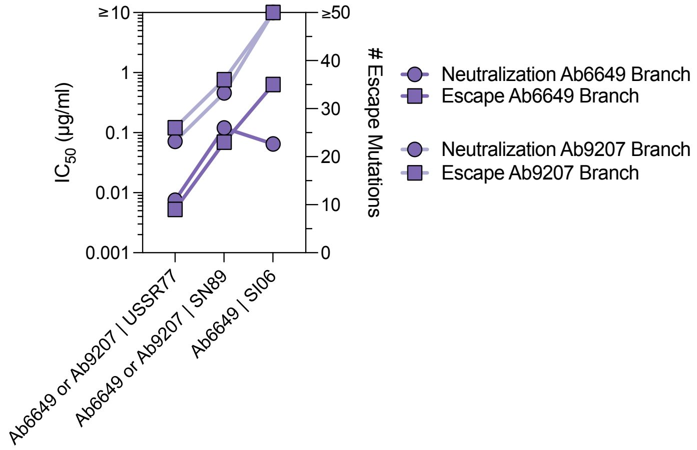


**S**

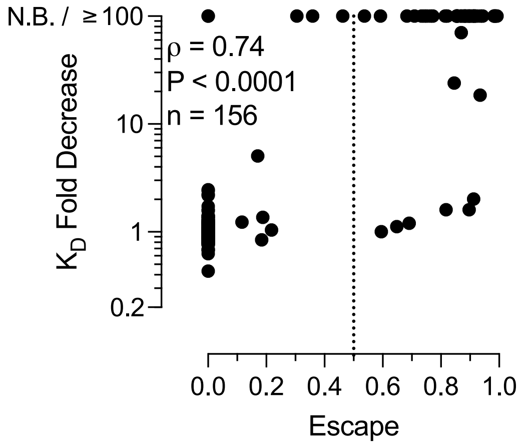


**T**

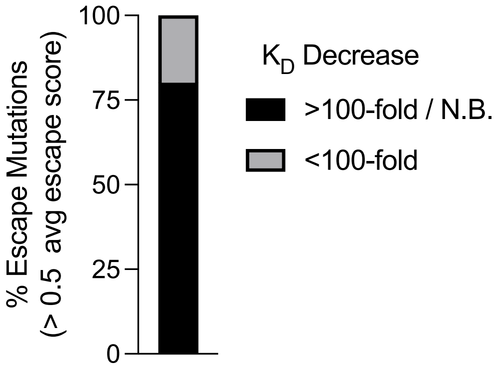


**Figure S3. Viral escape in drifted strains from affinity-matured antibodies. (A,G,M)** Neutralization potencies for lineages 860, 652, and 6649 against imprinting and drifted strains. Dotted lines indicate the concentrations used in selections. **(B,H,N)** Correlation between biological replicates for each mutation that shows positive differential selection in both replicates. Pearson's correlation coefficient is shown. Red box indicates our definition of escape mutations. The number inside the red box indicates the number of escape mutations. **(C,I,O)** Correlation between biological replicates summed by position. Pearson's correlation coefficient is shown. **(D,J,P)** Summed escape by position for all positions included in the libraries. All plots have a y-axis of 15. **(E,K,Q)** Heatmaps showing the relative escape for each individual selection, highlighting the positions that were selected by these antibodies. **(F,L,R)** Neutralization potencies and number of escape mutations shown on the same plot. **(S)** Correlation of avidities shown in Figure 3I with escape scores, which includes those data shown in Figure S2V. **(T)** Proportion of mutations with escape scores >0.5 (our definition of escape mutation) that have over >100-fold $K_D$ decrease from all escape selections.

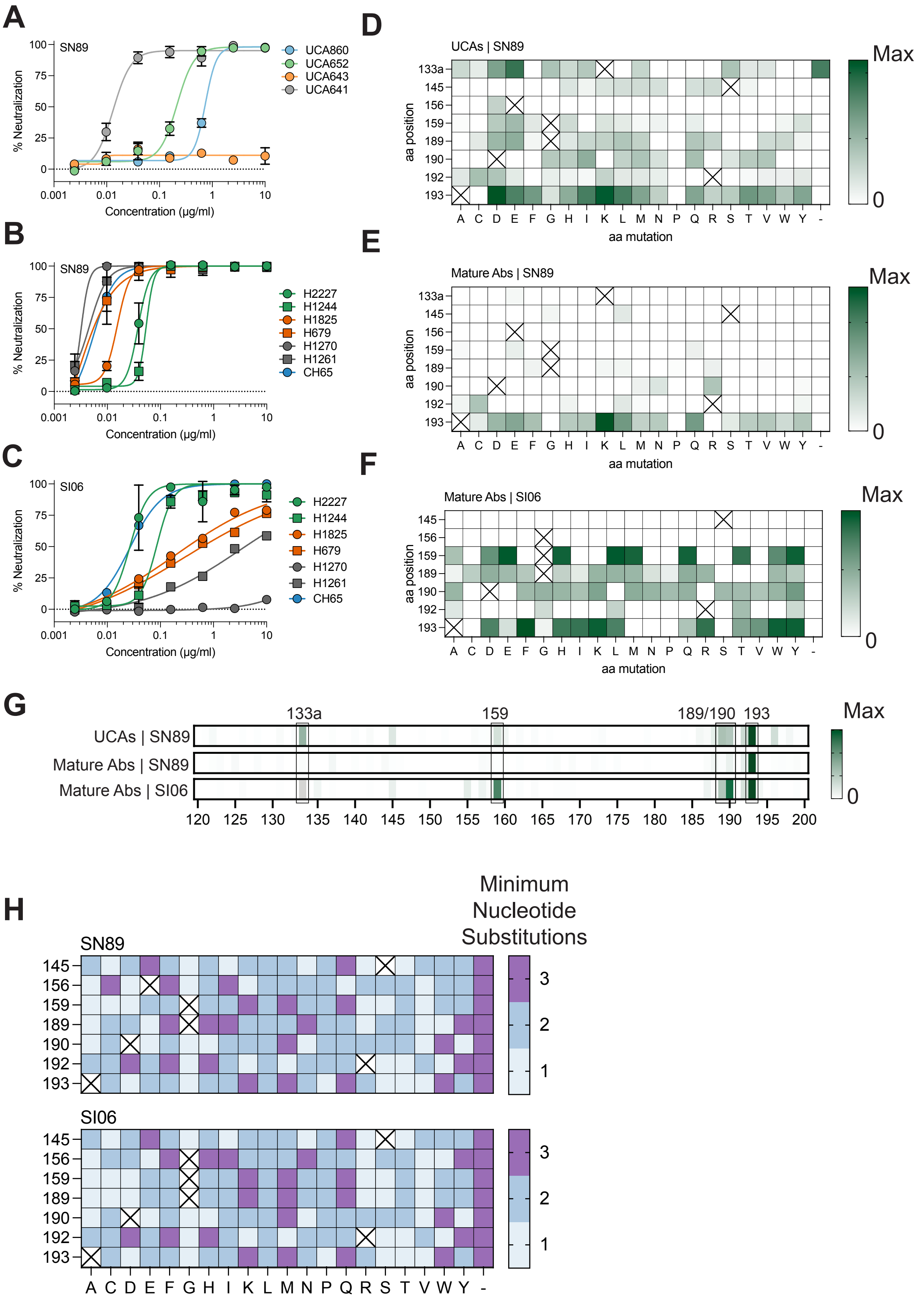


**Figure S4. Viral escape from an oligoclonal RBS antibody response. (A-C)** Neutralization potencies of the UCAs against SN89 **(A)**, the mature antibodies against SN89 **(B)**, and the mature antibodies against SI06 **(C)**. **(D-F)** Highly selected escape mutations. Each heatmap shows the summed effects of multiple antibodies (one per lineage) against a particular strain: UCA860, UCA652, UCA641 against SN89 **(D)**, CH65, H2227, H679, H1270 against SN89 **(E)**, and CH65 and H2227 against SI06 **(F)**. **(G)** Highly selected positions summed as in D-F. **(H)** The minimum number of nucleotide substitutions to acquire a specific mutation in SN89 (top) or SI06 (bottom).

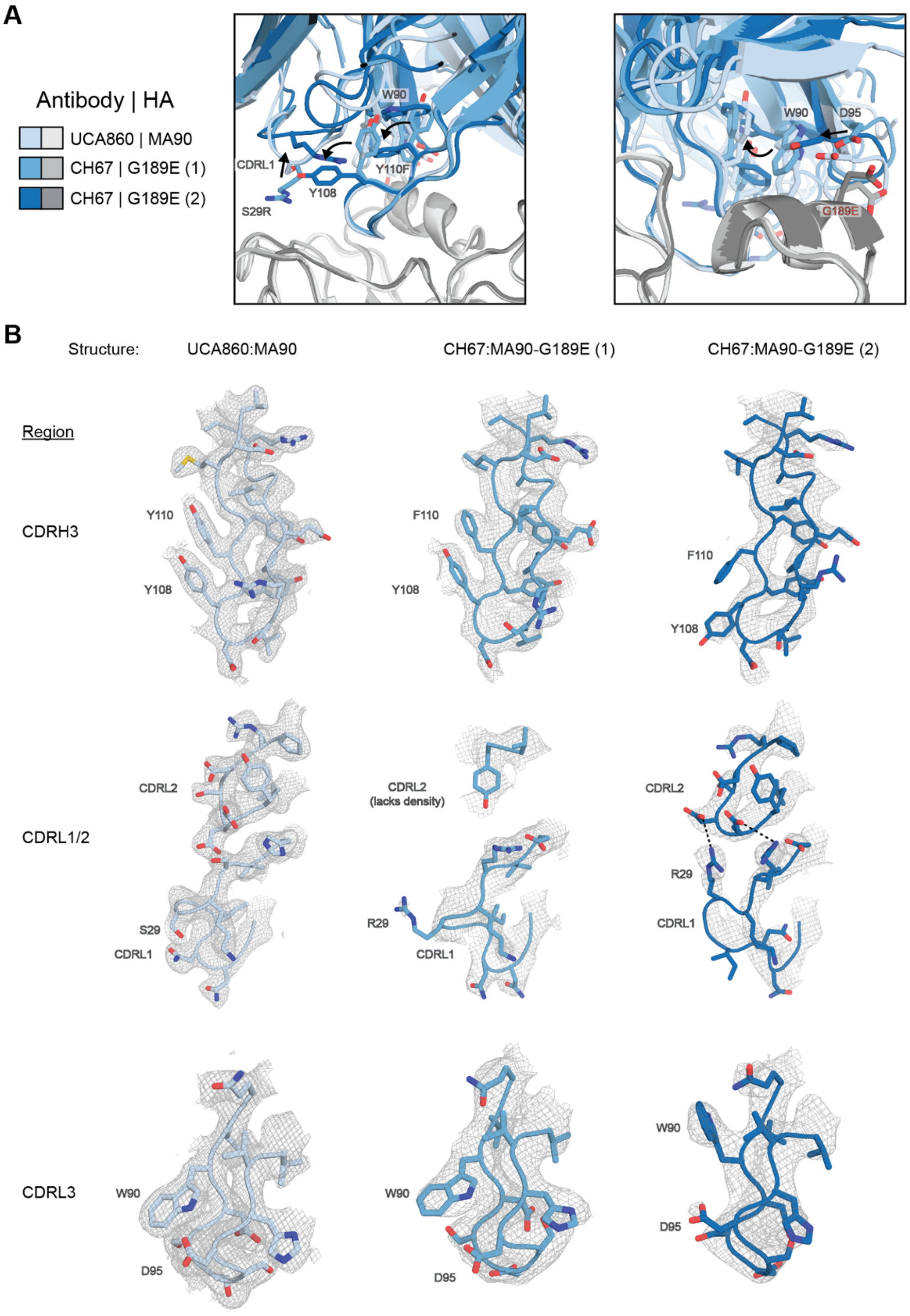


**Figure S5. Different structural conformations accommodate the G189E escape mutation. (A)** Superimposition of the UCA860|MA90 complex model with the CH67|MA90-G189E complex models. Residues shown adopt different conformations between structures. **(B)** 2Fo-Fc maps for each complex for the CDRH3, CDRL1, CDRL2, and CDRL3. Maps are contoured at 1σ and carved at 2Å.
